# Glomerular Elastic Modulus and Gene Expression Patterns Define Two Phases of Alport Nephropathy

**DOI:** 10.1101/2024.02.26.582201

**Authors:** Joonho Yoon, Zhenan Liu, Mathew Alaba, Leslie A. Bruggeman, Paul A. Janmey, Carlos A. Arana, Oluwatoyosi Ayenuyo, Isabela Medeiros, Sean Eddy, Matthias Kretzler, Joel M. Henderson, Viji Nair, Abhijit S. Naik, Audrey N. Chang, R. Tyler Miller

**Affiliations:** Department of Internal Medicine, University of Texas Southwestern Medical Center, Dallas, Texas, USA; Department of Internal Medicine, University of Michigan, Ann Arbor, Michigan, USA; Department of Inflammation & Immunity, Lerner Research Institute, Cleveland Clinic, Cleveland, Ohio, USA; Department of Physiology and Institute for Medicine and Engineering, University of Pennsylvania, Philadelphia, PA, 19104, USA; Department of Immunology, The University of Texas Southwestern Medical Center, Dallas, Texas, 75390, USA; Department of Pathology and Laboratory Medicine, Boston University Chobanian & Avedisian School of Medicine and Boston Medical Center, Boston, MA, USA; Pak Center for Mineral Metabolism and Clinical Research, UTSW Medical Center, Dallas, Texas, USA; Medicine Service, VA North Texas Health Care System, Dallas, TX, USA

**Author notes:** Corresponding authors- R. Tyler Miller, UT. Southwestern Nephrology, 8856, 5323 Harry Hines Blvd, Dallas, TX 75390. Abhijit S. Naik, Nephrology, University of Michigan, Department of Internal Medicine, Division of Nephrology, 1500 E Medical Center Dr, Fl 1, Ann Arbor, MI 48109. Co-first authors (Joonho Yoon, Zhenan Liu, Mathew Alaba). Co-senior authors (Abhijit S. Naik, Audrey N. Chang, R. Tyler Miller). These authors contributed equally to this work. The authors have declared that no conflict of interest exists.

## Abstract

Alport syndrome (AS), caused by COL4A3,4,5 mutations, leads to progressive glomerular disease and eventual kidney failure. In Col4α3^−/−^ mice (C57BL/6 background), we found that increased glomerular capillary deformability (reduced Young’s modulus, E) appears 2–3 months before detectable proteinuria or elevated serum creatinine. This early change indicates that podocyte injury precedes traditional clinical markers of disease and corresponds to reduced podocyte adhesion and loss. Bulk and podocyte-enriched RNA-seq data, obtained from deconvoluting bulk RNA sequencing data, showed that starting at 4 months, endoplasmic reticulum (ER) stress and the unfolded protein response (UPR) steadily rise while extracellular matrix remodeling, inflammation, epithelial–mesenchymal transition, and maladaptive repair begin. By 7 months, pathology shifts toward widespread parenchymal fibrosis, interleukin, cytokine, and chemokine signaling, cytoskeleton disruption, metabolic failure, and podocyte dedifferentiation. Notably, administration of the chemical chaperone Tauro-Urso-Deoxcycholic Acid (TUDCA) from weaning reduced ER stress, preserved glomerular stiffness, minimized podocyte detachment and loss, normalized inflammatory, injury, and fibrotic gene expression, and halved proteinuria and serum creatinine at 7 months, preserving kidney structure. Differentially expressed podocyte enriched genes from 4-month Col4α3^−/−^ (vs WT) mice were mapped to human orthologs in the NEPTUNE cohort. Four genes (*CRB2, GPC6, NKD1, STX11*) were associated with End Stage Renal Disease or 40% loss of eGFR (ESRD40). Collectively, our results identify these four genes and ER stress/UPR as key and treatable drivers of podocyte injury and disease progression in Alport syndrome, well before overt proteinuria, and highlight UPR activation and several genes as promising targets for disease-modifying therapies.

## Introduction

Alport nephropathy, caused by COL4A3,4,5 mutations, and other glomerular diseases, including Focal Sclerosing Glomerulonephritis (FSGS), and Minimal Change Disease (MCD), begin with podocyte injury that is followed by varying degrees of proteinuria, hematuria, and tubulo-interstitial damage. The first evidence of podocyte injury is disruption of foot process architecture, with effacement, flattening, spreading, and loss of slit diaphragms. Further injury leads to podocyte detachment and loss in the urine, and at some point, irreversible glomerular damage, tubulo-interstitial fibrosis with progression to End Stage Renal Disease (ESRD). The mechanisms underlying early podocyte injury in Alport nephropathy and the associated transcriptomic changes remain poorly defined. However, processes responsible for CKD progression, particularly tubulo-interstitial fibrosis, are increasingly understood.

Podocytes’ specialized cytoskeletal structure allows them to withstand traction and shear forces, thereby creating a specific biophysical environment for glomerular capillaries(1–4). Podocyte structure and function are maintained by factors including attachment to the Glomerular Basement Membrane (GBM), lateral attachment of foot processes to foot processes of neighboring podocytes, Ca+^2^ fluxes, various signaling pathways, and mechanical environment (stiffness of the GBM, shear forces, and stretch)(1, 2, 5–11). In rats, 5/6^th^ renal reduction causes glomerular capillary dilation to prevent increased shear stress(12). In acute and chronic glomerular injury models, including Alport nephropathy, protamine nephropathy, HIV nephropathy, ischemia, ischemia-reperfusion injury, and hypertrophy, glomerular capillary deformability increases as a result of podocyte injury(1, 2, 13–16). Intravital imaging studies show distension of glomerular capillaries and aneurysms in glomeruli from *Col4αa5^-/-^*mice at an early stage of disease, indicating increased deformability and reduced structural integrity of these capillaries(17–19).

The Alport GBM could injure podocytes through abnormal biochemical or biophysical differences between the Alport and normal GBM(8, 20). Alternatively, podocytes that produce COL4α3, 4, and 5 could be injured by the accumulation of mutant, misfolded collagen molecules, leading to activation of the unfolded protein response (UPR, ER stress) and additional cell-specific remodeling pathways(21–24). Additionally, abnormal glomerular capillary mechanics resulting from a disordered cytoskeletal structure could contribute to progressive podocyte injury by increasing susceptibility to mechanical forces. To begin to distinguish among and refine these injury mechanisms, we treated one group of *Col4α3^-/-^* mice with TauroUrsoDeoxyCholic Acid (TUDCA), a chemical chaperone that protects cells from UPR/ER stress-mediated injury or death(25–28). We analyzed renal function, glomerular capillary stiffness (E), structure, and renal RNA expression over the eight months of disease and mapped thos transcriptomic data to a different mouse model and human Nephrotic Syndrome Study Network (NEPTUNE) databases.

## Results

### Increased glomerular capillary deformability precedes proteinuria and increased creatinine in *Col4α3^-/-^*mice

We studied the *Col4α3^-/-^* mutation in the C57/Bl6 background to characterize progression of Alport nephropathy over the course of the disease. The Col3α3 mutation in *Col4α3^-/-^*mice is deletion of the *Col4α3* NC1 domain, where post-translational folding initiates, so it is a severe mutation(29–31). We measured Serum Creatinine (SCr µM) (**Figure 1A**) and urine Albumin/Creatinine Ratio (ACR mg/g) (**Figure 1B**) beginning at 2 months of age. From 4 months, SCr increased but did not reach statistical significance until 6 months, with a maximum value of 61 µM at 8 months. Similarly, urine ACR increased from 0.82 at 3 months to 122 mg/g at 8 months, reaching statistical significance at 6 months. The SCr and urine ACR values in the control mice remained unchanged over 8 months.

**Figure 1.**
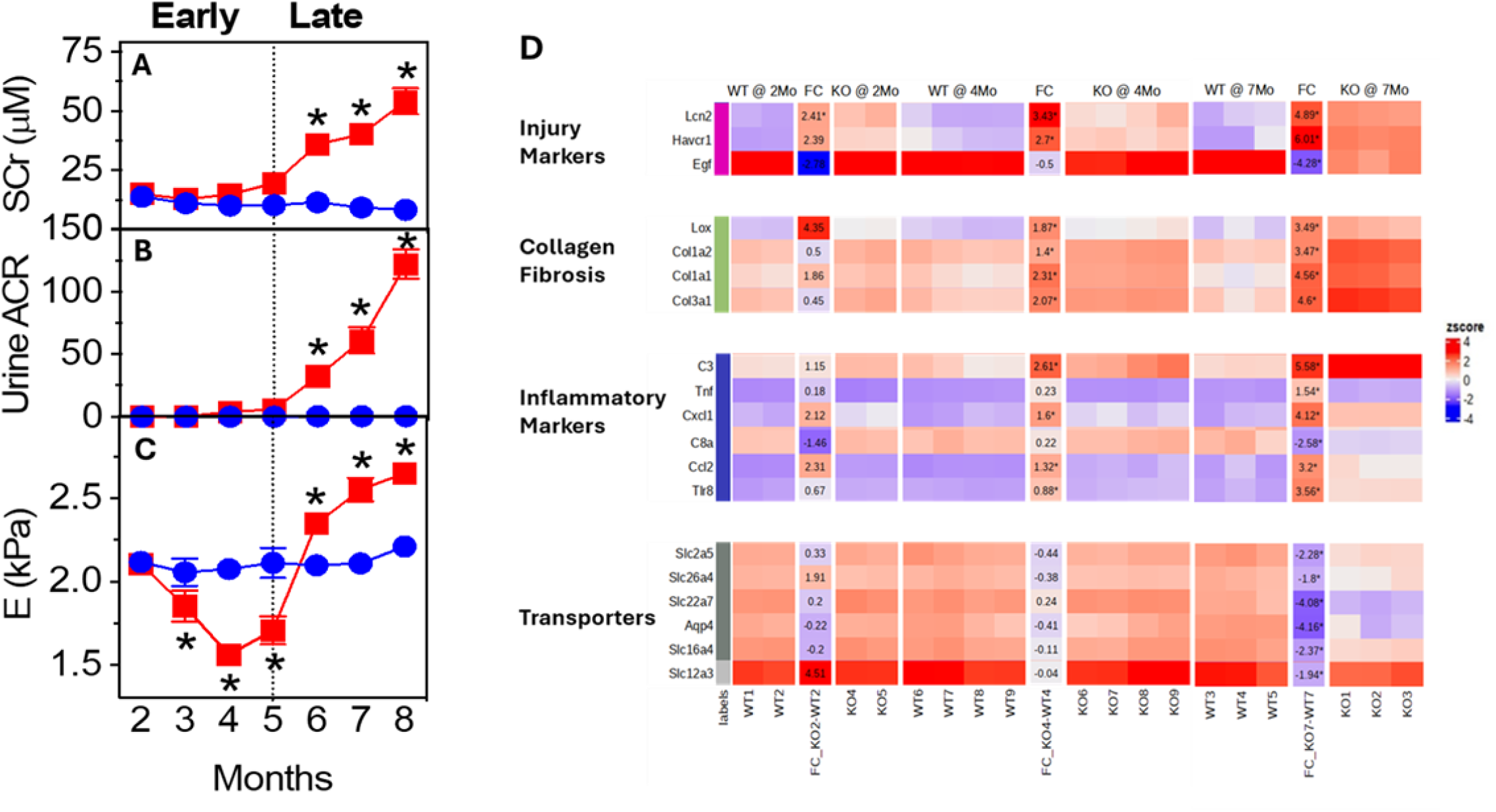
Measurement of SCr (**A**), urine ACR (**B**), and glomerular elastic modulus (E, Young’s modulus) (**C**) in Control (Blue) and *Col4α3^-/-^* (KO, red) mice over the course of renal disease compared to wildtype or heterozygote controls. Beginning two months after birth, SCr (µM) and urine ACR (mg/g) were measured monthly through eight months. Glomerular E (kPa) was measured in mice sacrificed at the indicated time points using microindentation of isolated glomeruli. Each point in the graphs shows average E values from 4 - 12 different mice with 6 – 10 glomeruli measured in each mouse. Significance was determined by 2-way ANOVA using Graphpad; **P<0.01, ****P<0.0001. **(D)** Differentially expressed genes in *Col4α3^-/-^* kidneys at 2, 4, and 7 months compared to age-matched WT. N=2 per group at 2 months, 4 per group at 4 months, and 3 per group at 7 months. Each column represents an individual mouse.

Previous work using kidney disease models demonstrated that glomerular capillary stiffness reflects podocyte cytoskeletal structure, and that with podocyte injury, capillaries exhibit increased deformability(1, 2, 13, 15). We measured glomerular stiffness, Young’s modulus (E), in the control and *Col4α3^-/-^* mice from 2 to 8 months of age using micro-indentation (**Figure 1C**)(32). The values for E for both the control and *Col4a3^-/-^* mice were identical at 2 months (2.2±0.2 kPa), and the values for the control mice remained essentially unchanged through 8 months. However, in *Col4α3^-/-^*mice, glomerular E values decreased at 3 months (1.85±0.09 kPa), reaching a minimum at 4 months (1.56±0.3 kPa). E then increased to values significantly greater than control by 7 months, reaching a maximum at 8 months (2.7±0.1 kPa). These directionally opposite changes in stiffness early and late in disease signify distinctly different structural and functional properties of glomerular capillaries over time (1–4). Reduced glomerular E before significant increases in SCr or urine ACR indicates that podocyte injury precedes these two events.

To understand the molecular processes or pathways activated or inactivated in this Alport model, we performed bulk RNA sequencing of kidney cortical mRNA at 2, 4, and 7 months of age (**Table S1, S2, S3**). We identified 2 Differentially Expressed Genes (DEGs) in the 2-month, 1346 DEGs in the 4-month, and 10,475 DEGs in the 7-month WT - *Col4α3^-/-^* comparisons. The heatmaps shown in **Figures 1D and S1** display the expression of selected genes, grouped vertically by function, from 2 to 7 months. From 2 to 4 months, gene cluster expression patterns remain relatively stable, reflecting minimal changes in renal function and histology (see below). However, at 2 months, two proximal tubule injury markers, *Lcn2* and *Havcr1*, are upregulated, and remain so with their greatest increases at 7 months. Beginning at 4 months, corresponding to the nadir of glomerular E, when negligible tubulo-interstitial disease is evident histologically, and before the onset of significant proteinuria, increased expression of collagen, fibrosis, and inflammation-associated genes is observed, along with genes associated with injury and repair, including *Sox4* and *Sox9*. By 7 months, expression of these genes increases rather than decreases, whereas *Hnf4a* expression decreases rather than increasing as it does during recovery from injury(33–35). By 7 months, the predominant pattern is maladaptive/failed repair, characterized by cell injury, death, and fibrosis(33–35). *Egf*, a marker of tubule health, decreases only at 7 months, when *Pck1, Fbp1*, and multiple tubule transporters, including *Aqp4* and *Ncc,* are also downregulated, demonstrating loss of differentiated tubule metabolic and transport functions.

For a better understanding of gene expression changes in our model, we performed an unbiased analysis of bulk RNA-seq data setting a threshold of |x| ≥ 0.5 to allow only highly relevant biological pathways and minimize statistical noise. All genes had adjusted p-values <0.05. We used the Gene Ontology 2025 tool, to test for enrichment of biological pathways, cellular processes, and molecular functions within the input gene set(36). **Table 1** provides supplemental table numbers that complement the supplementary figures across the manuscript. At 4 months, a total of 796 upregulated and 12 downregulated genes met this threshold. Top upregulated pathways included extracellular matrix (ECM), collagen fibril organization, and increased cell matrix adhesion. Increased responses to injury were reflected in genes associated with wound healing, epithelial-mesenchymal transition (EMT), and inflammatory responses, characterized by upregulation of phagocytic activity and cytokine receptor-mediated signaling (**Figure S2**, left panel; **Table S4)**. Pathways associated with MAPK, PI3K-Akt and Jak-STAT signaling were also upregulated. The increased expression of podocyte-specific markers, including *Nphs1, Nphs2, and Synpo,* was consistent with increased podocyte differentiation and hypertrophy. Increased expression of genes associated with Hippo-YAP-TAZ signaling (e.g., Tead3) provides evidence of increased podocyte differentiation and an ongoing compensatory response to podocyte injury and loss. The top-downregulated pathways, based on the 12 downregulated genes, are associated with basement membrane component synthesis and angiogenesis.

**Table 1.** Roadmap for supplemental tables and corresponding supplementary figures**, DEG:** Differentially expressed genes; **PE -** Podocyte Enriched genes**; WT:** Wild Type**; ACEi:** Angiotensin Converting Enzyme inhibitors.

|  | **Type of analysis** | **Corresponding Supplementary figures** |
| --- | --- | --- |
| **Table S1** | **2 mo** Col4a3<sup>-/-</sup> mice vs WT (Bulk RNA-seq) DEGs |  |
| **Table S2** | **4 mo** Col4a3<sup>-/-</sup> mice vs WT (Bulk RNA-seq) DEGs |  |
| **Table S3** | **7 mo** Col4a3<sup>-/-</sup> mice vs WT (Bulk RNA-seq) DEGs |  |
| **Table S4** | **4 mo** Col4a3<sup>-/-</sup> mice vs WT (Bulk RNA-seq) Pathway analysis | **Figure S2** |
| **Table S5** | Genes identified in ICA analysis in NEPTUNE and corresponding mouse orthologs | **Figure S3** |
| **Table S6** | **4 mo** Col4a3<sup>-/-</sup> mice vs WT (PE) DEGs |  |
| **Table S7** | **4 mo** Col4a3<sup>-/-</sup> mice vs WT (PE) Pathway analysis |  |
| **Table S8** | **7 mo** Col4a3<sup>-/-</sup> mice vs WT (Bulk RNA-seq) Pathway analysis | **Figure S4, Figure S5** |
| **Table S9** | **7 mo** Col4a3<sup>-/-</sup> mice vs WT (PE) DEGs |  |
| **Table S10** | **7 mo** Col4a3<sup>-/-</sup> mice vs WT (PE) Pathway analysis | **Figure S4, Figure S5** |
| **Table S11** | Morphometric analysis of glomerular lesions in light microscopic sections |  |
| **Table S12** | Semi-quantitative morphometric analysis of key features of glomerular injury from transmission electron microscopy images |  |
| **Table S13** | **4 mo** Col4a3<sup>-/-</sup> -T mice vs Col4a3<sup>-/-</sup> (Bulk RNA-seq) DEGs |  |
| **Table S14** | **4 mo** Col4a3<sup>-/-</sup> -T mice vs Col4a3<sup>-/-</sup> (Bulk RNA-seq) Pathway analysis | **Figure S9, Figure S10** |
| **Table S15** | Comparison of 60 podocyte enriched genes between Col4a3<sup>-/-</sup> mice vs WT over time and with TUDCA treatment | **Figure S11** |
| **Table S16** | DEGs from anti-Mir21 comparison in Rubel et al (Ref 48) |  |
| **Table S17** | DEGs from ACEi comparison in Rubel paper (Ref 48) |  |
| **Table S18** | Expression of mRNA coding for interleukins, cytokines and chemokines at 2, 4, and 7 months |  |
| **Table S19** | Primer sequences for qRT-PCR in Table 2 (Methods). |  |

Podocytes account for only 1% of the total cells in the kidney, so molecular signals from other cells in bulk RNAseq can overwhelm podocyte gene expression profiles. To obtain a precise podocyte response signal, we utilized micro-dissected bulk RNA-seq data of 234 glomerular profiles from the NEPTUNE study. An independent component analysis (ICA) identified a gene module enriched for podocyte-specific genes (n=327) in single-nucleus RNA-seq data from 77 NEPTUNE patients with a primary diagnosis of focal segmental glomerulosclerosis (FSGS)(37). The gene module identified thus represents podocyte-enriched genes deconvoluted from the bulk RNA-seq data. We then mapped these 327 podocyte-enriched genes to bulk RNA-seq data from all mice, identifying 291 mouse orthologs **(Figure S3A, Table S5)**.

We deconvoluted 4-month mouse bulk RNA-seq data to obtain a more podocyte-specific gene set and compared DEGs between *Col4α3^-/-^*and WT mice. Sixty genes were differentially expressed (**Table S6**). Using the same filtering threshold of |x| ≥ 0.5, a total of 50 genes were identified, 47 of which were upregulated and 3 were downregulated. Similar to the bulk RNA-seq data, the top upregulated processes included podocyte differentiation, EMT, and inflammation (**Figure S3B, Table S7**). The inflammatory responses were characterized by increased macrophage recruitment, consistent with the bulk data, and cellular responses to chemokines. Other upregulated pathways relate to podocyte cytoskeleton, including regulation of the actomyosin structural complex, stress fiber assembly, and filamin binding (**Figure S2, right panel)**. The 3 downregulated genes were those related to angiogenesis and the basement membrane (**Table S2**). The bulk RNA-seq data and the deconvoluted RNA-seq data thus complement each other. The bulk data enriched the transcriptional profile of all cells, while our deconvolution strategies identified several pathways known to be associated with podocytes, thereby confirming greater specificity.

Among the 5364 genes upregulated in the 7-month bulk data, as in the 4-month data, several pathways related to ECM organization, cell-matrix adhesion, and EMT were upregulated (**Figure S4, left panel, Table S8).** Increased cytokine activity, including Interleukin-6 (IL-6), IL-8, IL-10, IL-12, and TNF-mediated signaling was evident. Increased signal transduction activity was observed, including increased MAPK, PI3K-Akt, and Jak-STAT signaling. Pathways associated with cellular differentiation, migration, motility, cytoskeletal reorganization and cellular migration were upregulated. Of the 1119 downregulated genes, the top downregulated biological processes include oxidative phosphorylation, amino acid catabolic process, fatty acid beta-oxidation, branched-chain amino acid processes, sodium ion transport, gluconeogenesis, and the TCA cycle (**Figure S5, left panel, Table S8**).

We deconvoluted 7-month bulk RNA-seq data using the set of podocyte-enriched genes (**Table S9**), identified 168 DEGs in the *Col4α3*^-/-^-WT comparison and found 117 upregulated and 29 downregulated genes that met the thresholding criteria. Upregulated biological pathways were primarily associated with regulation of cytoskeletal dynamics, including cytoskeletal organization, stress fiber assembly, and actin-based movement, alongside processes linked to cellular differentiation and signaling, such as kidney development, regulation of epithelial-to-mesenchymal transition (EMT), neuron projection guidance, Wnt signaling pathway, and regulation of vascular endothelial growth factor (VEGF) production. Inflammatory responses were also enriched, including cytokine and chemokine production (**Figure S4, right panel, Table S10**). Additionally, pathways related to Golgi lumen function, which are associated with protein and lipid modification in the endoplasmic reticulum (ER), were upregulated. Downregulated pathways (**Figure S5, right panel, Tables S1A, S1B, S10**) included processes related to the regulation of cell proliferation, MAPK, and fibroblast growth factor receptor signaling. Based on transcriptomic data, there was also evidence of increased apoptosis and podocyte dedifferentiation, as indicated by reduced *Nphs1* expression. Lastly, the podocin (*Nphs2*) to nephrin (*Nphs1*) ratio (PNR) was elevated (2.45 vs. 1.37, p<0.05). An increased PNR reflects podocyte hypertrophic stress in studies of podocyte depletion and is linked to increased urinary podocyte detachment rates, proteinuria, and eventual kidney failure(38, 39).

### Correspondence between human and mouse inflammation-related gene expression patterns

Given the observed increase in inflammatory signaling and *Tnf* expression at 7 months, we tested whether *Col4α3^-/-^*mice showed a progressive increase in Tnf-mediated signaling compared with WT mice. We used the approach of Mariani et al. which showed increased expression of TNF-α signaling pathway genes in patients with FSGS and MCD, and defined a TNF activation gene signature (272 genes) and an activation score(40). To assess Tnf activity in our mice at all time points, we identified 205 mouse orthologs that matched the human TNF signature. **Figure 2** shows heatmaps depicting the 50 most upregulated and 50 most downregulated genes in the Tnf signature at 2, 4, and 7 months in *Col4α3^-/-^* mice compared to WT and shows a heatmap representing the activation score at each time point based on analysis of each mouse with calculation of a Z score. The activation score at 7 months in *Col4α3^-/-^* mice is significantly different from that of 7-month WT mice and *Col4α3^-/-^* mice at 2 and 4 months. However, other comparisons are not significant. *Tnf*, like many other inflammation-associated genes, was not differentially expressed at 2 and 4 months. These data indicate that not only does the Alport nephropathy mouse model express genes associated with proteinuria and poor outcomes in human renal biopsies of patients with glomerular disease, but also that there is an overlap in inflammatory TNF signaling between human and mouse datasets. Furthermore, in the *Col4α3^-/-^* mice, the magnitude of the change correlates with the stage of the disease.

**Figure 2.**
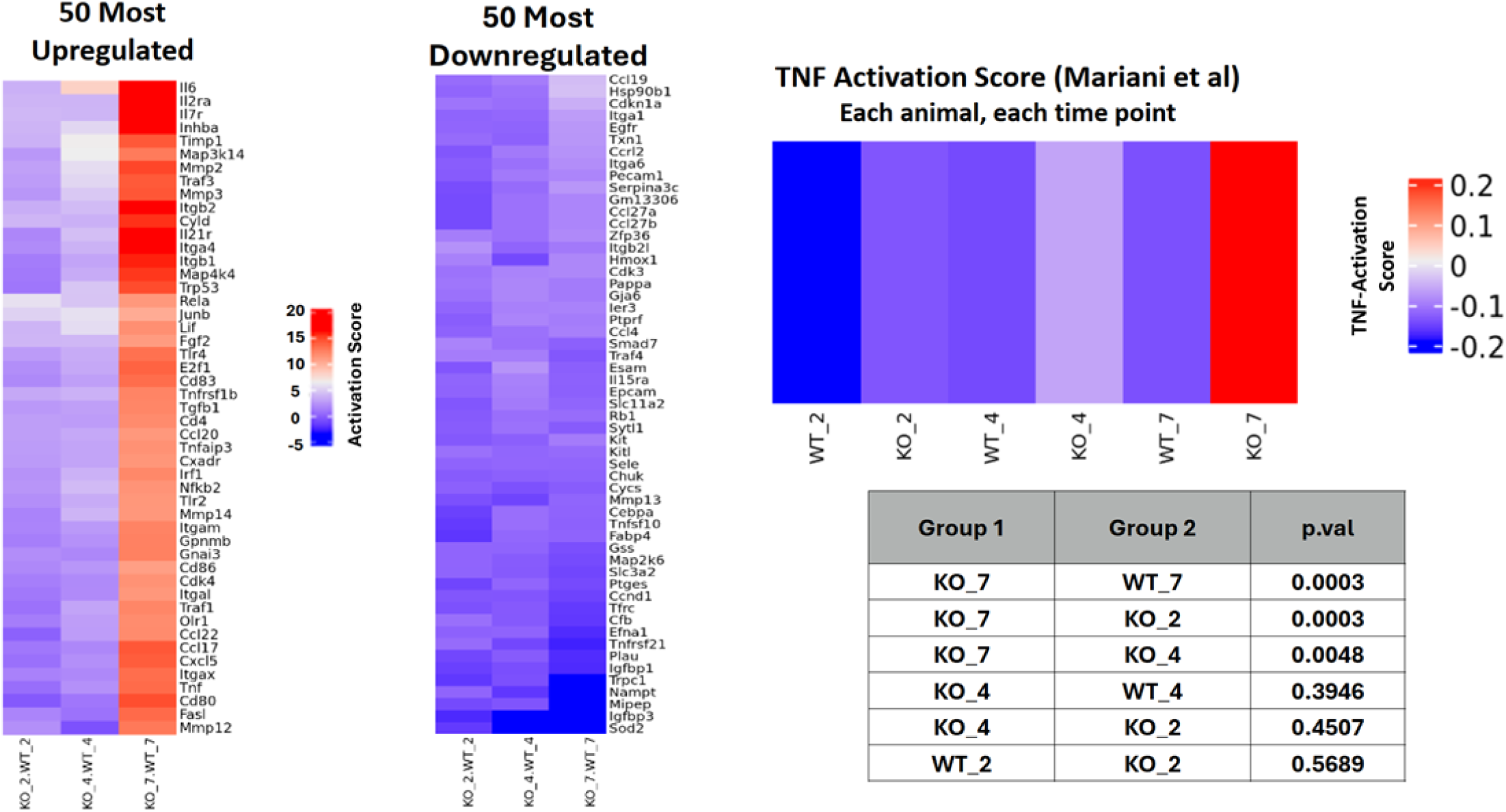
Genes and Gene Products Activated by TNF in Col4α3^-/-^ Mice Compared to NEPTUNE FSGS/MCD database. RNAseq data from WT and *Col4α3^-/-^*mice were analyzed for TNF signaling using the human 272 gene TNF network from Mariani et al(30). Mouse orthologs (205) of the 272-gene human TNF signature were identified. Individual gene expression values for each mouse were Z-transformed, and group mean gene expression was calculated. A) heatmaps showing the 50 most upregulated and 50 most downregulated genes. Each column corresponds to comparison of WT and *Col4α3^-/-^* mice at each time point (2 Mo - 2 mice WT and *Col4α3^-/-^*; 4 Mo – 4 mice WT and *Col4α3^-/-^*; 7 Mo – 3 mice WT and *Col4α3^-/-^*. The TNF activation score for each group of mice was the average Z-score of the 205 genes. B) Heatmap showing WT, *Col4α3^-/-^*, 2 – 7 Mo with significance of comparisons below.

### Evidence for unfolded protein pathway activation in *Col4α3^-/-^*podocytes

Podocytes provide mechanical support for glomerular capillaries, determine their elastic properties, form the slit diaphragms, and produce and secrete the Col4α3,4,5 heterotrimer for the GBM(41). Loss-of-function mutations in any one of these collagen chains cause reduced or failed heterotrimer secretion, leading to activation of the UPR in podocytes(24, 38, 42). Cells compensate for excess or misfolded protein by upregulating pathways that improve folding and/or protein degradation and reduce translation. We imaged WT and *Col4α3^-/-^* mouse cortex with confocal microscopy and found increased Bip (Hspa5) expression in glomerular podocytes (**Figure 3A**). The levels of Bip and Xbp1s protein and mRNA are increased in conditionally immortalized *Col4α3^-/-^* podocytes compared to their WT counterparts (**Figure 3B**). Cultured WT podocytes increase the expression of Bip, Xbp1s, and Ddit3 (CHOP, pro-apoptotic), in response to Tunicamycin, an agent that induces UPR, and expression of these genes is reduced in a dose-dependent manner by treatment with the chemical chaperone, TUDCA (23, 24, 26) (**Figure 3D**). Three genes related to UPR activation were differentially expressed in the *Col4α3^-/-^* - *Col4α3^-/-^*-T comparison: *Xbp1* (increased 1.85 X), *Dnajb9* (increased 1.52 X), and *Eef1akmt3* (decreased to 0.38 X). For these reasons and the expanded ER in *Col4α3^-/-^* glomerular podocytes (**Figure S6**), we conclude that an early step in podocyte injury is UPR activation, with disruption of normal protein processing(21, 22, 24).

**Figure 3.**
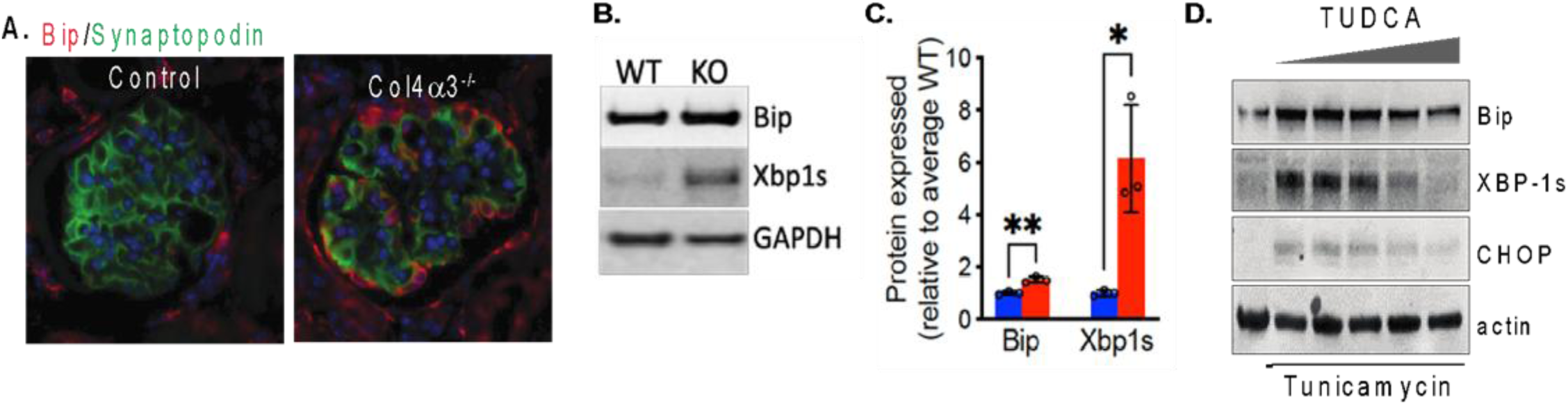
Evidence of UPR activation in glomeruli and podocytes from *Col4α3^-/-^* mice. **A)** Immunofluorescence images of Bip and Synaptopodin staining in 4-month WT and *Col4α3^-/-^* mouse glomeruli, **B)** Blots of UPR markers in podocyte cell lines under basal conditions derived from WT and *Col4α3^-/-^* (KO) mice. The cells were maintained under standard conditions, harvested in RIPA buffer at 70% confluence, and the crude extract was immunoblotted with antibodies against Bip, Xbp1s, and GAPDH. The conditionally immortalized WT and *Col4α3^-/-^* cells were developed from primary cultures of 3-month WT and Col4α3^-/-^ glomeruli. **C)** Comparison of Bip and Xbp1s protein expression from B (N=3, normalized to WT). **D)** Efficacy of TUDCA for blocking tunicamycin-induction of UPR marker expression in WT podocytes. Podocytes were treated with tunicamycin 1 µg/ml and then increasing concentrations of TUDCA from 1 - 4 mM for four hours.

Bulk-RNA-seq transcriptomic analysis over time showed progressive activation of the unfolded protein response (UPR) during endoplasmic reticulum (ER) stress(43). At 4 months, *Atf3* was significantly expressed, indicating early activation of the UPR through the PERK pathway. At the same time, genes like *Srebf1*, *Traf2*, and *Nfkb2* indicated IRE1-mediated stress signaling, immune responses, and activation of MAPK pathways (**Table S2**). By 7 months, a distinct UPR signature developed. Genes including *Atf3, Atf6, Atf6b, Ddit3, Eif2ak3, Ern1, Hspa5, and Xbp1* were upregulated (**Table S3**), encoding key UPR transducers like IRE1α, PERK, and ATF6, and effectors such as BiP and XBP1s. This pattern reflects increasing ER stress adaptation, including translational attenuation, chaperone induction, and ER-associated degradation.

### TUDCA treatment slows progression of *Col4α3^-/-^* nephropathy

The role of ER stress in Alport nephropathy was investigated by feeding *Col4α3^-/-^* mice TUDCA incorporated into mouse chow (0.8%) from the time of weaning, 21 - 25 days old, through the end of studies and comparing physiologic, morphologic, and transcriptomic changes among WT, *Col4α3^-/-^*, and TUDCA-treated *Col4α3^-/-^* mice (Col4α3^-/-^-T) at 4 and 7 months(44). At 4 months, SCr and urine ACR values were similar among all three groups (**Figure 4A and B**, left top two panels). However, at 7 months, average ACR and SCr values were respectively 90±11 mg/g and 55±7 µm in the Col4α3^-/-^ mice; values for SCr and ACR in the Col4α3^-/-^-T mice were 40±8.6 mg/g and 33±3.9 µm (**Figure 4A A and B**, right top two panels). At 4 months, the value for E in the WT mice was 2.1±0.04 kPa, 1.56±0.05 kPa for the *Col4α3^-/-^* mice, and 2.0±1.0 kPa in the *Col4α3^-/-^*-T mice (**Figure 4C**). At 7 months, WT glomerular E was 2.1±0.06 kPa, and for the *Col4α3^-/-^* glomeruli it was 2.70±0.05 kPa, while the value for the *Col4α3^-/-^*-T glomeruli was 1.9±0.04 kPa, a value comparable to WT.

**Figure 4.**
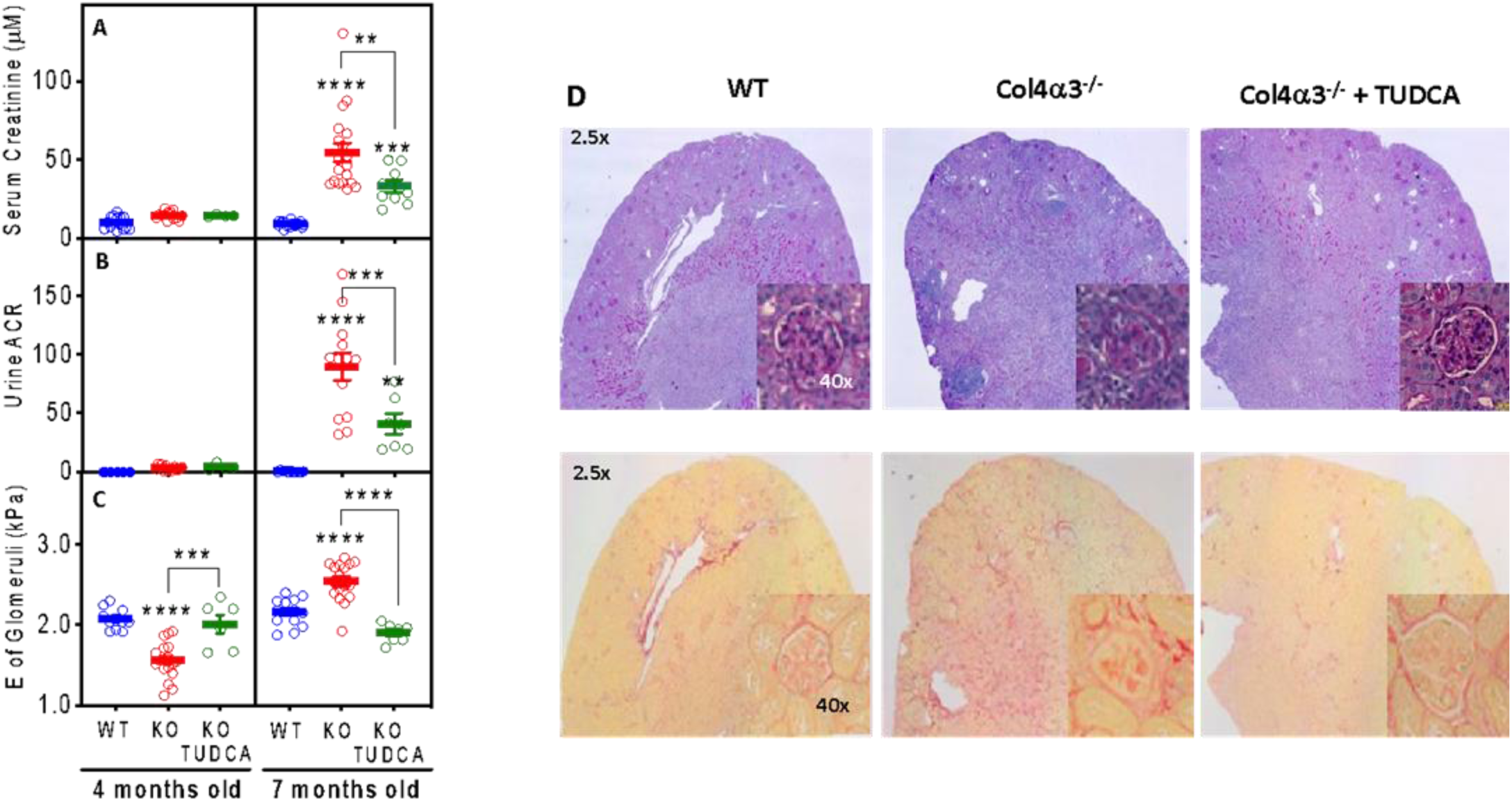
TUDCA treatment slows the progression of renal disease and minimizes changes in E in Col4α3^-/-^ mice. Col4α3^-/-^ (KO) mice were treated from weaning (approximately 4 weeks) with 0.8% TUDCA mouse chow (KO TUDCA) until sacrifice. At 4 and 7 months of age, SCr (µM) (**2A**), urine ACR (**2B**), and glomerular elastic modulus (E, kPa) (**2C**) was measured in WT (Blue), KO (Red) and TUDCA-treated KO animals (Green). In A and B, each circle represents a measurement from one mouse, and in C, each circle represents E measured from an average of 6-10 individual glomeruli from a single mouse. Significance was determined by ANOVA and multiple comparisons test using GraphPad; **P<0.01, ***P<0.001, ****P<0.0001. **2D.** Comparison of kidney cortex from WT, *Col4α3^-/-^* (KO), and TUDCA treated *Col4α3^-/-^* (KO) animals. Top row, sections from 7-month mice are stained with PAS and bottom row sections from 7-month mice are stained with picrosirius red.

### TUDCA treatment preserves *Col4α3^-/-^* kidney structure

Chronic kidney disease is marked by progressive fibrosis and loss of renal function, a component of which can be attributed to proteinuria(45). Our physiologic measurements at 7 months (**Figure 4A-C**) showed preserved renal function in the *Col4α3^-/-^*-T mice, implying preserved glomerular and renal structure. Low magnification kidney sections from 4-month mouse kidneys stained with PAS or Picrosirius Red (PSR) show that sections from WT, *Col4α3^-/-^*, and *Col4α3^-/-^*-T mice are indistinguishable (**Figure S8**), indicating that at the 4-month time point, overall renal parenchymal injury is minimal. At seven months (**Figure 4D**), the extent of injury in the *Col4α3^-/-^* kidneys is evident in the PAS and PSR-stained sections. The PAS-stained *Col4α3^-/-^* kidneys show fibrosis, inflammatory cell infiltrates, and disrupted renal architecture. In the *Col4α3^-/-^*-T kidneys, these features are present, but to a significantly lesser extent. The insets show representative images of glomeruli from each group. The glomeruli from the *Col4α3^-/-^*mice are sclerotic and have excess, irregular PSR staining. Although glomeruli from the *Col4α3^-/-^*-T mice are not normal, their structure is relatively preserved compared to the untreated *Col4α3^-/-^*glomeruli.

Expression of genes associated with renal injury and fibrosis at 7 months was compared in WT and *Col4α3^-/-^*mice using qRT-PCR (**Table 2**). Expression of these genes increased in the *Col4α3^-/-^* mice compared to WT and was reduced to levels intermediate between WT and untreated *Col4α3^-/-^* mice by TUDCA. *Egf* mRNA, a marker for renal tubular health, decreased significantly in the *Col4α3^-/-^* mice to 20% of the WT value, but TUDCA restored its expression to 70% of WT. TUDCA slows the progression of nephropathy in *Col4α3^-/-^* mice, in part, by reducing cytokine and chemokine production and fibrosis.

**Table 2.** Summary of qRT-PCR of selected fibrosis, cytokine, chemokine, or injury-related genes in 7-month *Col4α3^-/-^* mice without (KO) and with (KO-TUDCA) TUDCA treatment, compared to age-matched WT littermates (100%). All values in KO and KO-TUDCA groups represent fold change from 4-month-old WT ± SD as determined by ANOVA and are significant to p<0.05 or better. The number of samples in each group is shown in parentheses after the values (N).

| **<u>Fibrosis</u>** | **<u>WT (N)</u>** | **<u>KO (N)</u>** | **<u>KO-TUDCA (N)</u>** |
| --- | --- | --- | --- |
| **Col1α1** | **1 (4)** | **55.5 ± 13 (3)** | **26.2± 6.2 (4)** |
| **Col1α2** | **1 (4)** | **29.6 ± 7.3 (5)** | **13.9 ± 3.2 (4)** |
| **Col3α1** | **1 (4)** | **74.1 ± 31.9 (5)** | **31.9 ± 2.4 (4)** |
| **Lox** | **1 (4)** | **29.7 ± 7.4 (3)** | **13..3 ± 7.6 (4)** |
| **Loxl2** | **1 (4)** | **21.5 ± 7.1 (3)** | **6.7 ± 3.1 (4)** |
| **<u>Cytokines</u>** |  |  |  |
| **TGF-β1** | **1 (4)** | **4.8 ± 0.8 (4)** | **3.0 ± 0.7 (4)** |
| **TNF-α** | **1 (4)** | **30.3 ± 7.9 (3)** | **14.3 ± 3.0 (4)** |
| **IGF-1** | **1 (4)** | **9.81 ± 2.23 (4)** | **5.93 ± 1.72 (4)** |
| **<u>Chemokines/Receptors</u>** |  |  |  |
| **Cxcl1** | **1 (4)** | **45.0 ± 11.4 (4)** | **20.9 ± 2.11 (3)** |
| **Ccl2** | **1 (4)** | **41.5 ± 9.3 (4)** | **34.6 ± 5.68 (4)** |
| **Ccr2** | **1 (4)** | **15.0 ± 3.45 (5)** | **4.04 ± 1.57 (3)** |
| **<u>Injury Markers</u>** |  |  |  |
| **EGF** | **1 (4)** | **0.20 ± 0.1 (5)** | **0.7 ± 0.2 (4)** |
| **Ccn1 (Cyr61)** | **1 (4)** | **4.46 ± 0.6 (3)** | **2.0 ± 1.0 (4)** |

### TUDCA preserves *Col4α3^-/-^* mouse podocyte number

Glomerular capillaries become deformable at 3 months, before significant increases in SCr or urine ACR, indicating that podocyte injury begins at or before that time. In Alport patients, podocytes are shed into the urine at an accelerated rate, and podocytes can be detected in urine before significant proteinuria develops(18, 19). We compared glomerular podocyte density in WT, *Col4α3^-/-^*, and *Col4α3^-/-^*-T mice at 2, 3, and 4 months using WT1 staining as a podocyte marker (**Figure 5**)(46). At 2 months, before the reduction of glomerular E in the *Col4α3^-/-^* mice, the podocyte density was similar among all three groups. At 3 months, when increased deformability was detectable in the *Col4α3^-/-^* glomeruli, podocyte density remained unchanged in the three groups and was comparable to the values for 2-month WT glomeruli. In the 3-month *Col4α3^-/-^* mice, the SCr and ACR values were similar to those for the WT mice. However, at 4 months, podocyte density was reduced by 40% in *Col4α3^-/-^*glomeruli, while TUDCA treatment protected glomeruli from podocyte loss.

**Figure 5.**
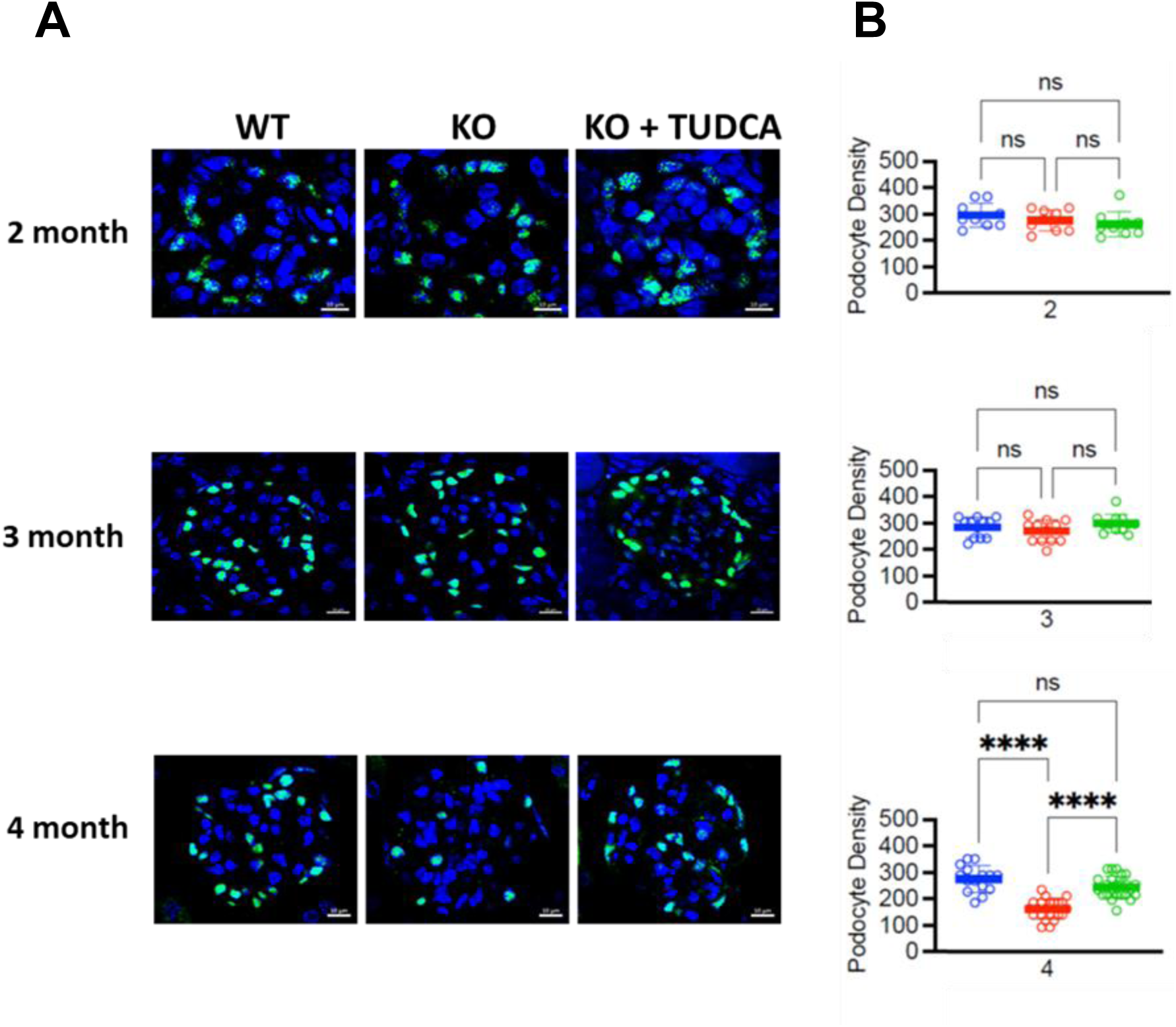
TUDCA reduces podocyte loss from *Col4α3^-/-^* mouse glomeruli. **A)** Representative glomeruli in kidney sections from WT, *Col4α3^-/-^* (KO), and TUDCA-treated *Col4α3^-/-^*(KO+TUDCA) mice from 2, 3, and 4- 4-month-old mice were stained for WT-1 and DAPI. Scale bar: 10 µm. **B)** Summary of podocyte density analysis. WT – Blue, *Col4α3^-/-^* - Red, and *Col4α3^-/-^* + TUDCA Green. The number of WT-1 and DAPI-positive nuclei in 20 glomeruli from 3 mice was counted and used for podocyte density calculations. Each dot represents the number of WT-1 positive nuclei in a single glomerulus. Data are shown as mean ± SEM and significance was determined by ordinary one-way ANOVA, followed by Dunnett’s multiple comparisons test, ****P<0.0001. At 2 months - Podocytes/10^6^ µm^3^ ± SD, WT 296±44, *Col4α3^-/-^* 276±39, and *Col4α3^-/-^*+TUDCA 262±45). At 3 months - Podocytes/10^6^ µm^3^ ± SD, WT 285±35, *Col4α3^-/-^* 270±40, and *Col4α3^-/-^*+T 299±32). At 4 months, Podocyte numbers were /10^6^ µm^3^ ± SD, WT 276±48, *Col4α3^-/-^*162±37, and *Col4α3^-/-^*+TUDCA 246±40).

### Reduced podocyte adhesion corresponds to early glomerular capillary deformability

We expected that podocyte injury should be detectable at 3 months as decreased adhesion, because capillaries are abnormally deformable and *Col4α3^-/-^*podocyte numbers are preserved. WT and *Col4α3^-/-^* glomeruli had similar patterns of synaptopodin and nephrin staining, indicating crudely preserved structure (**Figure S8**). We measured podocyte adhesion as podocyte loss *in vivo* following volume loading. Mice were injected with 0.5N saline IP, and urine was collected over 2 hours(47). Urinary podocytes were quantified using two methods: immunoblot analysis of urine for nephrin normalized for creatinine (**Figure 6A, B**), and counting WT-1-positive cells in the urine sediment (**Figure 6C**). Urinary nephrin increased fourfold over WT in the *Col4α3^-/-^* samples, and TUDCA treatment reduced it by approximately half. In the urine sediment, podocytes were present only in *Col4α3^-/-^* mice (**Figure 6C**). These observations were consistent across 3 experiments, in which, although the total number of podocytes lost in the urine varied from mouse to mouse, only the *Col4α3^-/-^* mice had multiple WT-1-positive cells in their urine.

**Figure 6.**
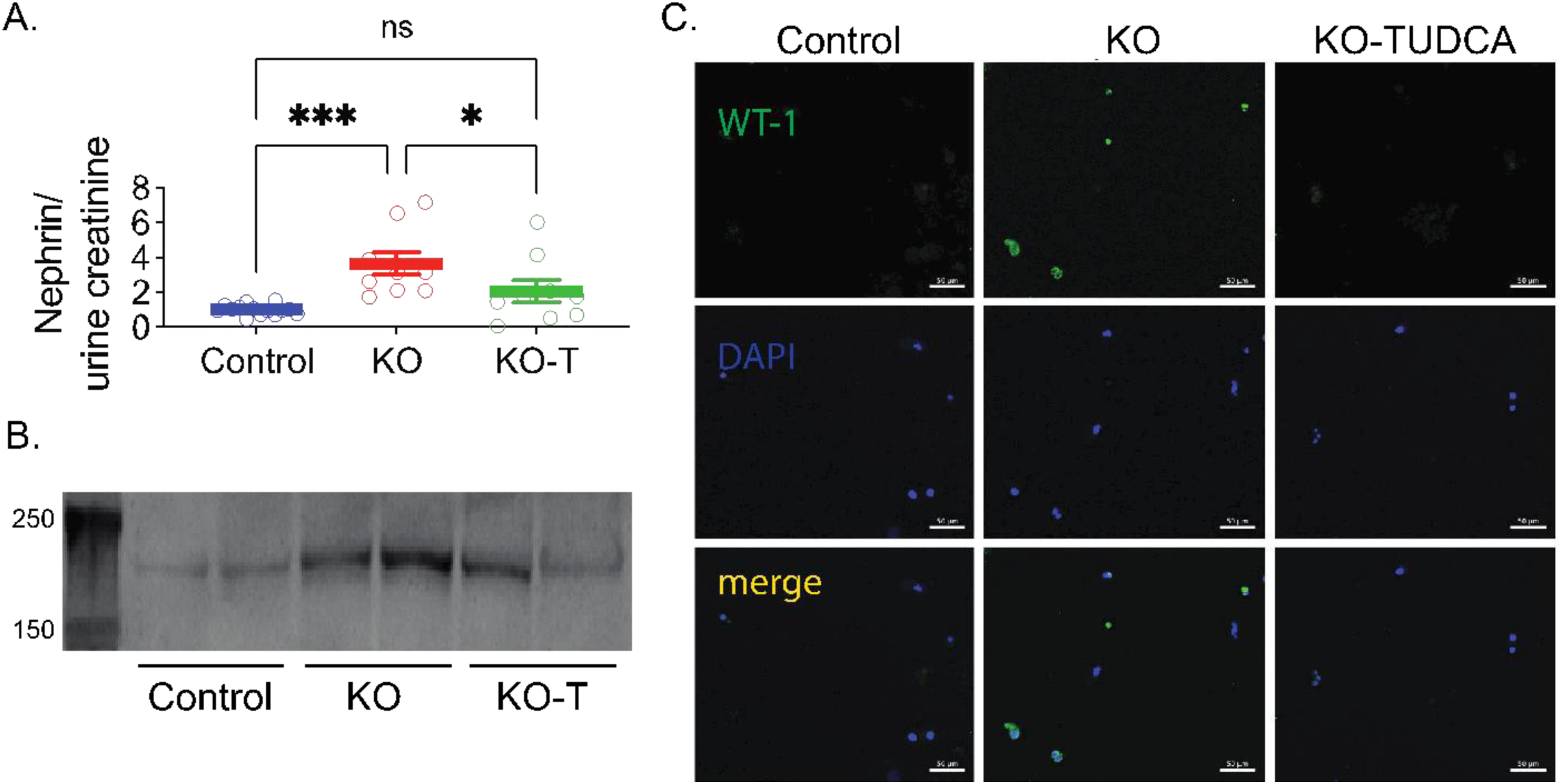
Glomerular capillary deformability and podocyte adhesion. WT (Control), *Col4α3^-/-^*(KO), and *Col4α3^-/-^* + TUDCA (KO-TUDCA) mice were injected (IP) with 0.5N saline (80 µl/g IP), and urine was collected over the subsequent two hours. The urine samples were normalized for creatinine, immunoblotted for nephrin, and quantified with densitometry. The urine pellet was stained for WT-1 and DAPI. **A**) Aggregate data for all sample groups expressed as nephrin/creatinine (arbitrary densitometry units for nephrin) for each sample group. WT – Blue, *Col4α3^-/-^* - Red, and *Col4α3^-/-^* + TUDCA Green. Each point represents a sample from a single mouse; the bar indicates mean±SEM. Significance was determined by ordinary one-way ANOVA, followed by Tukey’s multiple comparisons test; *P<0.05, ***P<0.001. **B**) Representative urine nephrin immunoblot with duplicate samples, loading normalized with creatinine. **C**) Representative urine sediment samples stained for WT-1 and DAPI.

### TUDCA protects glomerular capillary and podocyte structure

Light microscopic images of glomeruli from 4-month-old *Col4α3^-/-^*mice show generally preserved structure, but the glomeruli are hypertrophic compared to WT mice and exhibit multiple dilated capillaries (**Figure 7, top row**). Focal lesions are present, but there is little evidence of tubulo-interstitial damage. The *Col4α3^-/-^*-T glomeruli show evidence of focal lesions and capillary dilation, but considerably less than the *Col4α3^-/-^* glomeruli and are not hypertrophic. At 7 months, the patterns described above persist, but in the *Col4α3^-/-^*mice, they are more severe, characterized by increased glomerulosclerosis, focal glomerular lesions, and significant tubulo-interstitial injury and inflammation. The *Col4α3^-/-^*-T kidneys show less glomerular and tubulo-interstitial injury than the *Col4α3^-/-^*kidneys, but injury progressed from the 4-month time point (**Table S11).**

**Figure 7.**
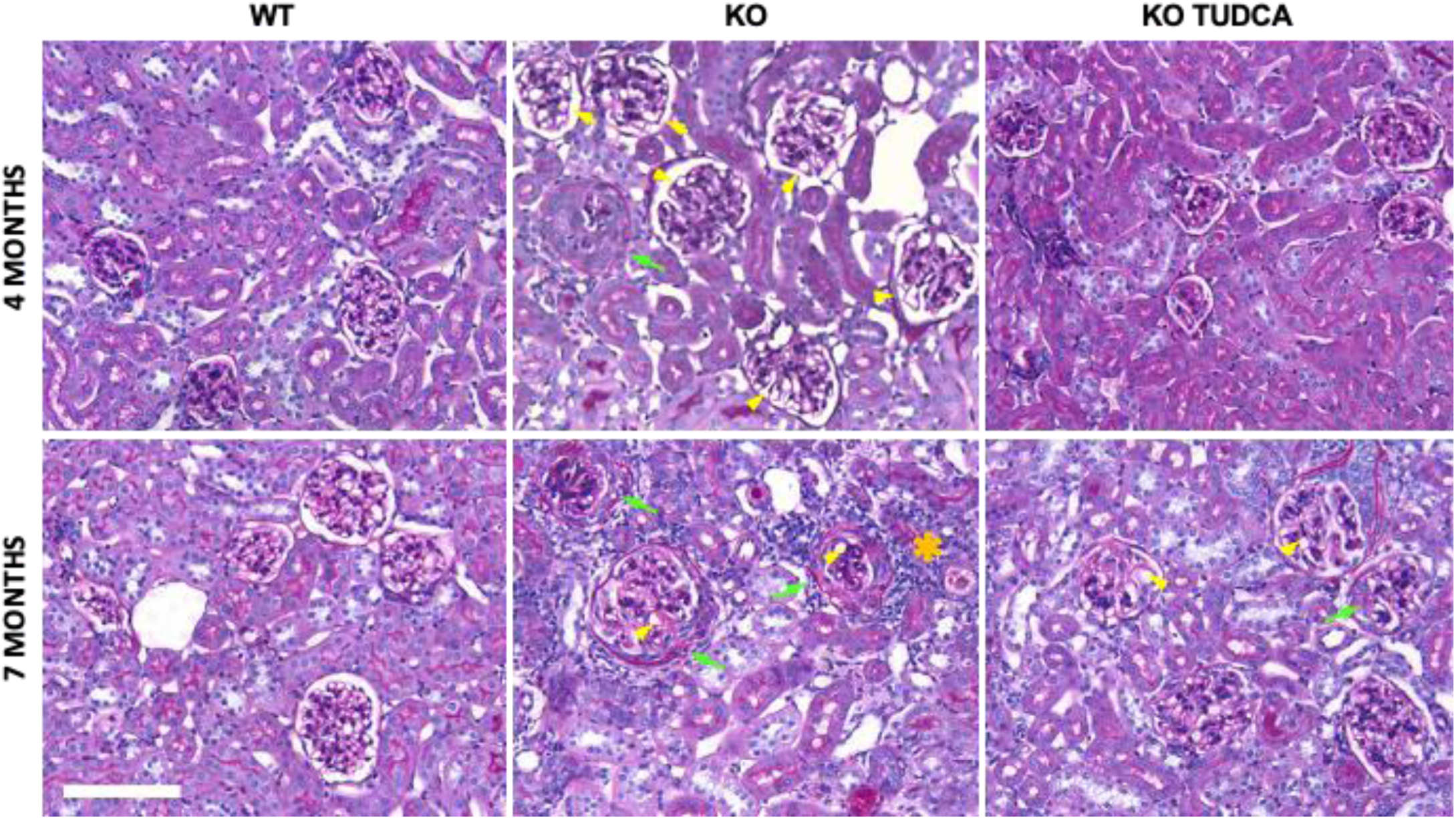
Photomicrographs of PAS-stained kidney sections from 4-month-old (top row) and 7-month-old (bottom row) WT, *Col4α3^-/-^* (KO), and TUDCA-treated *Col4α3^-/-^* (KO TUDCA) mice. The 4-month KO image shows glomerular enlargement, focal proliferative glomerular lesions (green arrow), and dilation of glomerular capillary lumens (yellow arrowheads) that are not observed in Control (top left) or the TUDCA-treated KO mice (top right). The 7-month KO mice (bottom center) show more extensive glomerular proliferative lesion development (green arrows), with proliferation composed predominantly of epithelial cells filling Bowman’s space, along with patchy areas of tubular atrophy and interstitial fibrosis with chronic inflammation (orange asterisk). The section from the 7-month TUDCA-treated mice shows evidence of less severe glomerular injury. Scale bar = 200 µm for all images.

We compared podocyte and GBM structures in our three groups of mice at 4 and 7 months using transmission electron microscopy (TEM)(**Figure 8**). At 4 months, the glomeruli from *Col4α3^-/^*^-^ mice exhibit regions of irregular GBM structure, characterized by both thin and thick regions. The podocyte foot processes are normal in some regions but also show evidence of effacement and widening in other regions. Some podocyte cell bodies appear hypertrophic, exhibit vacuolization, have expanded ER indicative of ER stress, and form microvilli (**Figure S6)**. In the *Col4α3^-/-^*-T glomeruli, these pathologic findings are present, but less severe than in the *Col4α3^-/-^* mice. By 7 months, the pathologic changes in the *Col4α3^-/-^*GBMs and podocytes are more advanced, with evidence of substantial glomerulosclerosis. The *Col4α3^-/-^*-T glomeruli have characteristics intermediate between the WT and Col4α3^-/-^ glomeruli. The pathologic findings are quantified in **Table S12**.

**Figure 8.**
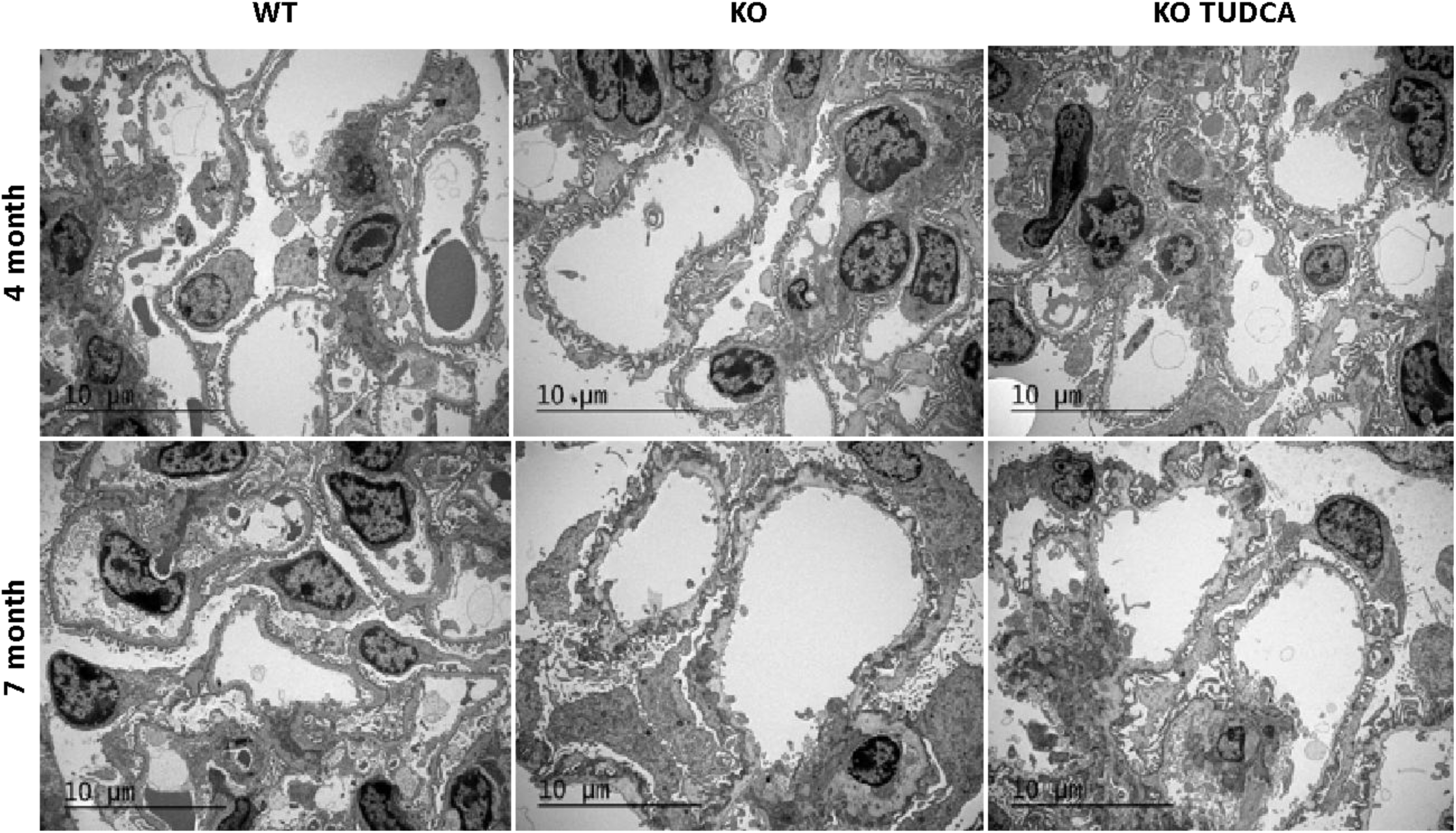
Transmission Electron micrographs comparing glomeruli from 4- (top row) and 7- (bottom row) month WT, *Col4α3^-/-^* (KO), and TUDCA-treated *Col4α3^-/-^* (KO TUDCA). Four-month glomeruli from KO mice (top center) and KO TUDCA mice (top right) have features of segmental podocyte injury, including mild segmental podocyte foot process effacement, more extensive but segmental foot process irregularity, and mild microvillous change, that are concurrent with segmentally prominent “knob-like” subepithelial basement redundancy, characteristic of the *Col4α3^-/-^* mouse model. WT mice (top left) show none of these features. TUDCA treatment reduces the severity and extent of glomerular pathology. Seven-month (bottom row), KO glomeruli (bottom center) reveal more severe and widespread podocyte injury, with extensive podocyte foot process effacement and irregularity, cellular hypertrophy, and microvillous change that are accompanied by GBM changes including extensive subepithelial basement redundancy, segmental “basket weave”, and segmental glomerular capillary loop dilation. In contrast, 7-month KO TUDCA mice (bottom right) show podocyte and glomerular basement membrane changes that are similar in nature to those of the KO mice but are less extensive and more segmental. None of these changes is seen in WT glomeruli (bottom left). Magnification – scale bar = 10 µm.

**Figure 9.**
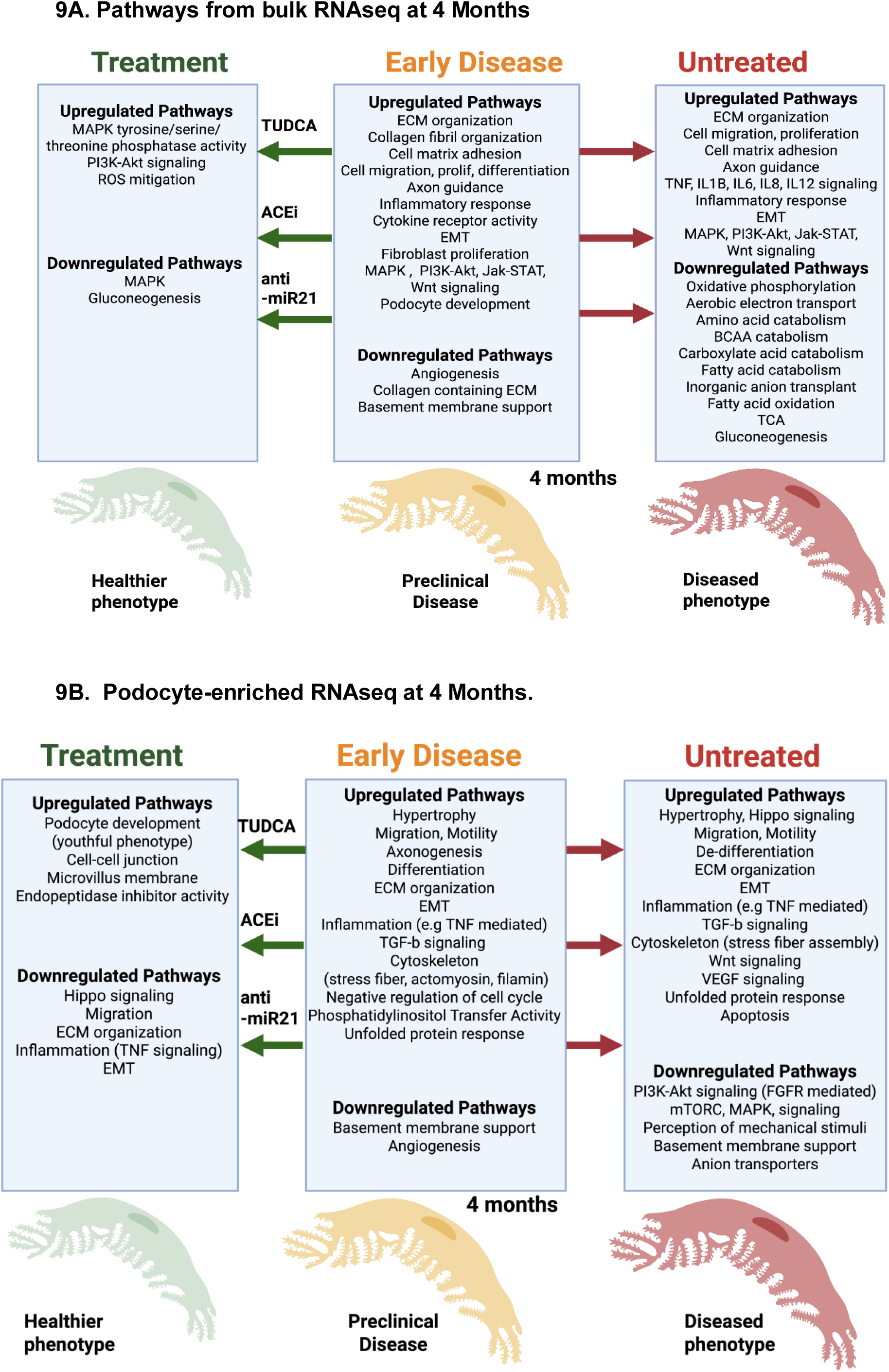
Summary integrating transcriptomic changes in treated (TUDCA, ACEi, anti-Mir21) vs. untreated 4 Mo Col4α3^-/-^ mice using 4 Mo bulk (9A) and PE (9B) data. The middle top panel shows the basic transcriptomic effects of Alport nephropathy on podocyte gene expression. TUDCA, anti-Mir21, and ACEI reduce these disease-dependent transcriptional effects to suppress the Alport disease phenotype.

### TUDCA normalizes the transcriptomic profile of 4-month-old *Col4α3^-/-^* mice

We assessed TUDCA effects on gene expression in 4-month-old *Col4α3^-/-^* mice using bulk RNA sequencing, as this age represents a critical transition point in disease progression. Using the same thresholding criteria as earlier, we identified 62 upregulated genes out of 128 DEGs (**Table S13**). The top enriched biological pathways were those that negatively regulate MAPK signaling, increase the activity of genes related to MAPK tyrosine/serine/threonine phosphatase activity, and Phosphatidylinositol 3-Kinase/Protein Kinase B Signal Transduction (PI3K/PKB) (**Figure S9, left pane**l; **Table S14**). Genes coding for proteins involved in the mitigation of Reactive Oxygen Species (ROS), *e.g., Gstm, Gstt1, Txnrd3,* were also upregulated. Gluconeogenesis was downregulated. Consistent with a reduction in MAPK signaling, cellular growth and proliferative responses were downregulated *(Ccnd1, Egfr)* **(Figure S10, left panel, Table S14).** These data suggest that at the bulk-RNA level, TUDCA affects signaling and metabolism, reducing MAPK signaling, promoting anti-apoptotic PI3K/PKB activity, and normalizing cellular metabolism, growth, and proliferation, while increasing the generation of ROS-scavenging proteins, all resulting in preservation of phenotype and kidney function.

We evaluated the effect of TUDCA on the directionality of the 60 genes (**Table S6**) that were differentially expressed between *Col4α3^-/-^*-T mice untreated *Col4α3^-/-^* mice using the same deconvolution strategy described above to identify altered transcriptomic programs related to preservation of podocyte structure and function (**Figure S11**)(37). *Col4α3^-/-^*-T mice exhibited a nonsignificant reversal of expression in 26 genes and an increase in expression in 34 genes (**Table S15**). To refine our analysis, we restricted enrichment to genes with log2 fold changes ≥0.1 or ≤-0.1. Using this approach, 14 genes reversed their expression direction, and 19 genes increased their expression.

Pathway enrichment analysis of genes reversed with TUDCA revealed suppressed pathways related to podocyte development and differentiation (e.g., *Nphs1* and *Nphs2*), as well as those involved in basement membrane synthesis, all of which were upregulated in the Col4α3^-/-^ disease state (**Figure S9, right panel, Table S16**). In contrast, TUDCA enhanced pathways associated with reduced cell migration and angiogenesis. (**Figure S10, right panel**). Thus, TUDCA shifted disease-associated gene expression patterns toward the WT pattern.

Since the 60 podocyte-enriched genes were hypothesized to be causally related to Alport nephropathy progression, we investigated other models and analyzed a dataset from an F1 hybrid *Col4α3^−/−^* mouse model (129/SvJ × C57BL/6J) (**Figure S12**)(48). These mice were treated with either anti-miR-21 or angiotensin-converting enzyme inhibitors (ACEi), starting at 4 weeks of age, and were sacrificed at 15 weeks for RNA-seq analysis. Using our deconvolution approach, we extracted expression values for the 60 genes and compared the effects of anti-miR-21 or ACEi with TUDCA treatment(37). In the anti-miR-21-treated group, 49 of 60 genes were differentially expressed (**Table S17**), whereas in the ACEi-treated group, 38 of 60 genes were differentially expressed (**Table S18**). In the TUDCA-treated group, none of the genes were differentially expressed compared to the anti-miR-21 and ACEi groups. Despite the lack of a statistically significant difference in expression in the TUDCA group, the direction of the mRNA response to therapy was highly consistent across all three therapeutic interventions. The weaker response to TUDCA compared to anti-miR-21 and ACEi could be attributed to the advanced disease state (evidenced by proteinuria) already established by 4 weeks in the anti-miR-21 and ACEi-treated groups, resulting in a higher-amplitude signal for disease suppression.

### Differentially Expressed Genes in 4-month-old *Col4a3^-/-^* Mice Mapped to Human Orthologs Predict Long-Term Function in Glomerular Disease

To assess the relevance of the 60 genes identified in the 4-month podocyte-enriched data from *Col4a3^-/-^* mice to human health, we performed Cox regression to examine the association between these genes and clinical outcomes in NEPTUNE patients. Outcomes were defined as time to end-stage renal disease (ESRD) or a 40% decline in estimated glomerular filtration rate (eGFR) from baseline (ESRD40). To address multicollinearity in this high-dimensional dataset, we used a Lasso model, which identified a total of eight biologically relevant genes: *SLC14A2, NKD1, CRB2, GPC6, NPHS1, FOXC1, STX11* and FRY. and found that four, *NKD1, CRB2, GPC6*, and *STX11,* were associated with risk for ESRD40. In a multivariable Cox model adjusted for age, gender and race, a log2-fold increase in the expression of *CRB2* and *GPC6* was associated with hazard ratios (HRs) of 1.62 (95% CI: 1.20–2.20) and 2.62 (95% CI: 1.42–4.80) for the risk of kidney failure, respectively. Conversely, a log2-fold increase in *NKD1* expression was associated with an HR of 0.37 (95% CI: 0.15–0.90), and for *STX11,* an HR of 0.42 (95% CI 0.21 – 0.87) **(Figure S13).**

## Discussion

Our study provides a detailed, time-resolved characterization of Alport nephropathy progression in *Col4a3^-/-^* mice and identifies ER stress in podocytes as an early, therapeutically actionable driver of disease. Notably, we observed that increased glomerular capillary deformability, reflecting podocyte cytoskeletal dysfunction, precedes detectable proteinuria or a rise in serum creatinine by at least 2-3 months. This early biophysical alteration coincides with activation of UPR, reduced podocyte adhesion, and accelerated podocyte detachment, establishing a sequence in which proteotoxic stress initiates podocyte injury before conventional markers of kidney damage appear.

The biophysical measurements performed in our study demonstrate a biphasic change in glomerular capillary Young’s modulus: an initial softening (decreased E) at 3-4 months, followed by later stiffening consistent with progressive glomerulosclerosis and ECM accumulation. Our report buttresses findings from other studies that show that podocyte injury reduces capillary wall stiffness and extends this by demonstrating that the early biophysical failure is the earliest quantifiable abnormality in this Alport model, occurring even before podocyte loss(1, 2, 4). The preservation of podocyte number at 3-months, despite abnormal capillary biomechanics and increased urinary podocyte loss, is consistent with functional detachment or weakened adhesion rather than cell death as the primary mechanism of early biomechanical failure.

Transcriptomic profiling revealed two genes that were differentially expressed at 2 months. There was robust activation of injury, ECM, inflammatory, and EMT pathways by 4 months when capillary deformability reached its nadir. Deconvolution of bulk-RNA-seq data using a podocyte-enriched gene module confirmed that several of these early signals originate from podocytes, including upregulation of actomyosin stress fiber assembly, filamin binding, and Hippo-YAP/TAZ signaling, consistent with compensatory hypertrophy and dedifferentiation responses to mechanical and proteotoxic stress. The early upregulation of *Lcn2* and *Havcr1*, even before the presence of inflammation or tubulointerstitial damage, further supports the concept of primary glomerular injury causing secondary tubular injury.

Three major stress-responsive signaling cascades—MAPK, JAK-STAT, and PI3K-Akt—were among the earliest and most consistently activated pathways at 4 months and remained highly enriched at 7 months. MAPK signaling is a well-established downstream effector of ER stress via IRE1α–TRAF2–ASK1 and PERK–eIF2α–ATF4 axes, and its sustained activation drives podocyte EMT, actin remodeling, and apoptosis while promoting fibroblast activation and ECM deposition(49). JAK-STAT, particularly STAT3, is activated by multiple cytokines that are induced secondary to ER stress and initial podocyte injury; persistent STAT3 signaling in podocytes and tubular cells perpetuates inflammation and fibrosis(50). PI3K-Akt activation, also evident from 4 months onward, reflects compensatory survival signaling in stressed podocytes but paradoxically contributes to hypertrophy, autophagy suppression, and eventual dedifferentiation(51–53). The simultaneous engagement of these three interconnected pathways creates a self-reinforcing maladaptive network that amplifies early proteotoxic injury into progressive glomerulosclerosis.

A central mechanistic insight is the early and progressive activation of the UPR in *Col4a3^-/-^* podocytes. Because the loss of the NC1 trimerization domain leads to the retention of misfolded collagen IV α3 chains in the ER, this induces BiP overexpression and XBP1 splicing causing subsequent activation of the PERK and ATF6 branches. Because of prolonged ER stress, multiple podocyte functions are likely impaired including deficits in the synthesis and maintenance of the slit diaphragm, cytoskeletal integrity, and adhesion to the glomerular basement membrane (GBM). As observed on electron microscopy, features such as expanded ER, vacuolization, and microvillus formation demonstrate this proteotoxic burden. Importantly, because UPR activation occurs before inflammatory escalation and fibrosis, ER stress acts as an upstream initiator rather than a consequence of later-stage inflammation.

The therapeutic efficacy of the chemical chaperone TUDCA provides additional strong evidence for the pathogenic role of ER stress. Initiated at weaning, TUDCA completely prevented the early decrease in glomerular stiffness, largely preserved podocyte adhesion and number, markedly reduced proteinuria and serum creatinine at 7 months, and attenuated glomerulosclerosis, inflammation, and fibrosis. At the transcriptomic level, TUDCA normalized disease-associated pathways—suppressing MAPK hyperactivation (evidenced by upregulation of dual-specificity phosphatases), enhancing ROS detoxification, and shifting podocyte-enriched gene expression toward wild-type patterns. The partial rather than complete protection likely reflects the fact that TUDCA addresses proteotoxic stress incompletely and indicates that primary or initiating injury mechanism may not be the GBM defect or the mechanical consequences of the absence of α3α4α5(IV) heterotrimers. Comparison with previously published data in a faster-progressing 129×B6 F1 *Col4a3^-/-^*model further contextualizes TUDCA’s effects: both anti-miR-21 (which directly represses MAPK/ERK and TGF-β signaling) and ACE inhibition (which reduces angiotensin II–driven JAK-STAT and MAPK activation) produced stronger reversal of the same 60-gene podocyte signature than TUDCA, consistent with their later initiation in already proteinuric mice and their direct targeting of downstream fibrogenic pathways(48). The fact that TUDCA nonetheless shifted the same genes in a similar direction—despite being given prophylactically in a slower-progressing strain—underscores that alleviating upstream ER stress is sufficient to intercept the entire maladaptive MAPK/JAK-STAT/PI3K cascade before it becomes established.

Of translational relevance, several of the 60 podocyte-enriched genes dysregulated at the critical 4-month transition point in mice have orthologs that predict progression to kidney failure in human glomerular disease (NEPTUNE cohort). Notably, *CRB2* and *GPC6* upregulation and *NKD1* and *STX11* downregulation in humans were independently associated with ESRD40. Three of these 4 genes (*CRB2, GPC6, NKD1*) were expressed at higher levels in podocytes than other renal cell types supporting the validity of bulk data deconvolution (**Figure S14**). The expression of *STX11* is minimal across all cell types. STX11 is involved in vesicular trafficking and regulating protein transport among late endosomes and the trans-Golgi network, but it has not been associated with kidney disease(54). The reasons for the relative lack of expression of *STX11* in the single nuclear RNA-sequencing data remain unclear but could be the result of lower sequencing depth. However, its presence in this group of genes and response to TUDCA, anti-miR-21, ACEi suggest that it may have a role in podocyte injury, possibly slit diaphragm protein trafficking. CRB*2* is a key component of the slit diaphragm. It forms a protein complex with other molecules (e.g., *NEPH1*, NPHS1) that maintains the structure of the slit diaphragm and podocyte foot processes, and functions as a dynamic sensor to recognize other Crb2-expressing cells triggering increased adhesion(55, 56). *CRB2* is highly expressed in podocytes relative to other renal cell types (**Figure S14**). Overexpression studies (e.g., transgenic mice) show that excess *Crb2* disrupts the balance of the Crumbs complex, leading to disrupted slit diaphragm structure, foot process effacement, and proteinuria. *CRB2* upregulation is observed in human collapsing glomerulopathy and some forms of FSGS, and it may function as a modifier gene in Alport syndrome(57). Although *GPC6* is highly expressed in podocytes, it has not been studied in that setting, while *GPC5* which is linked evolutionarily and structurally has been. Both *GPC5* and *GPC6* are cell-surface heparan sulfate proteoglycans and participate in Wnt signaling. We found upregulation of *GPC6* with Alport nephropathy, its association with ESRD40, and its downregulation by TUDCA(58). Increased podocyte *GPC5* expression is associated with human nephrotic syndrome and its knockdown is protective(59). Podocyte specific knockdown of *Gpc5* in diabetic Akita mice preserves the expression of nephrin and attenuates foot process effacement suggesting its role in maintaining the slit diaphragm(60). *GPC6* stimulates intestinal elongation via hedgehog and non-canonical Wnt pathways, so although its function in podocytes is not established, it is likely to be important(61). *NKD1* is expressed in podocytes and is a negative regulator of canonical Wnt signaling(62). Transient Wnt/β-catenin signaling in the adult is important for tissue repair, but persistent expression leads to fibrosis and disrupted slit diaphragm structure(63). Reduced *NKD1* expression as seen in our data could perpetuate Wnt/β-catenin signaling and contribute to ESRD40. Because these 4 genes were identified among the 60 genes that were differentially expressed in the 4-month podocyte-enriched data from *Col4a3^-/-^* mice (vs. WT), this 4-gene signature is relevant in a primary podocytopathy like Alport syndrome. These data suggest that the 4-gene signature may forecast ESRD or loss of kidney function years before it is evident clinically providing a valuable diagnostic tool for clinicians.

In addition to increased Wnt signaling, we found evidence of persistent activation of the MAPK and Jak-Stat pathways in our model. These are developmental and/or fundamental regulatory pathways that are important for repair and normal cell function, but genes like *Sox4* and *Sox9* adversely affect repair and function when activated persistently(33–35, 37, 64). This positioning not only reinforces the fidelity of the *Col4α3^−/−^* model but also suggests that early podocyte stress signatures may serve as biomarkers of future progression in patients with Alport syndrome or other proteinuric glomerulopathies.

We observed that TNF-mediated signaling escalates dramatically between 4 and 7 months, mirroring patterns seen in human FSGS associated with poor outcomes, suggesting a feed-forward loop in later stages of the disease, in which podocyte ER stress drives cytokine/chemokine release, leading to an inflammatory response that drives fibrosis, injury, and potentially an irreversible disease stage. Many inflammatory mediators follow this pattern of late appearance in the bulk RNAseq data (**Tables S18A and B**). Mitigating ER stress early in the course of disease with TUDCA preserved podocyte number, reduced proteinuria and blunted this inflammatory surge. This result suggests that upstream targeting of podocyte proteotoxic stress that reduces proteinuria that is tubulo-toxic may be more effective than downstream anti-inflammatory or antifibrotic strategies alone(45).

In conclusion, our work is consistent with ER stress-induced podocyte dysfunction as the primary event in Alport nephropathy, manifesting first as altered glomerular biomechanics and podocyte detachment, then activating downstream pathways that drive inflammation, fibrosis, and irreversible glomerular and tubulo-interstitial scarring. TUDCA slows disease progression when started early, providing a strong rationale for clinical trials of chemical chaperones or other UPR-modulating agents in children and young adults with Alport syndrome, ideally before proteinuria develops.

## Methods

### Reagents

Fluorescent antibodies were purchased from Invitrogen. Primary *Antibodies for immunofluorescence* **-** Bip: ab21685, Abcam; WT-1: sc-7385, Santa Cruz; Synaptopodin: sc-21537, Santa Cruz; Nephrin: AF3169, R*&*D Systems. *Antibodies for immunoblotting* - Bip: ab21685, Abcam; Xbp1s: sc-7160, Cell signaling, and 24868-1-AP, Proteintech; Chop: (D46F1), #5554, Cell Signaling; GAPDH: XP, #5174S, Cell signaling; β-actin – #AM4302 Invitrogen.

### Mouse breeding and treatment

Col4α3^-/-^ mice (C57Bl/6J) were obtained from Jeff Miner(36) with genotyping by Transnetyx. Homozygous males were bred with heterozygous females to obtain Col4α3^-/-^ and Col4α3^+/+^ mice. Col4α3^-/-^ mice were fed 0.8% (3α,7β-Dihydroxy-5β-cholanoyltaurine, TUDCA incorporated into mouse chow from the time of weaning (25-28 days old) until sacrifice(25, 26, 28, 30). TUDCA (98% purity) was from Chemimpex and incorporated into Teklad standard mouse chow. SCr was measured by submandibular puncture or exsanguination at sacrifice by standard methods. UACR was measured using spot urine and standard methods.

Podocyte adhesion was measured in vivo by injecting 0.5N saline (80 µl/g IP) and collecting urine over two hours(47). An aliquot of urine was normalized for creatinine and immunoblotted for nephrin. Another aliquot was centrifuged, and the sediment on a slide was stained for DAPI and WT-1. Mouse glomeruli were isolated by sequential sieving as previously described and maintained in DMEM with 0.1% FBS at room temperature before stiffness measurements or imaging(1, 2, 65).

### Podocyte cell line development and culture

Podocyte cell lines were derived from WT and Col4α3^-/-^ primary glomerular podocyte outgrowths isolated from adult mice (12-16 weeks of age). Glomeruli were maintained in DMEM with 0.1% FBS at room temperature and seeded onto collagen I-coated dishes for 4-6 days, trypsinized, and re-plated for experiments or immortalization. Primary podocytes were conditionally immortalized by infection with a lentivirus expressing the temperature-sensitive SV40 T antigen tsA58. Lentivirus was produced from pLenti-SV40-T-tsA58 (abm cat# LV629) as previously described(66) using the VSV envelope from pMD2.G (Addgene cat#12259), and the packaging plasmid pCMV delta R8.2 (Addgene cat # 12263), gifts from Didier Trono. Primary glomerular outgrowths were infected for 48 hours, followed by selection with puromycin (2ug/ml). Conditionally immortalized mouse podocytes were cultured in RPMI 1640 with 10% FBS. Cell cultures were expanded at 33°C and, for experiments, differentiated at 37°C for 7-10 days as previously described(67).

### Animal Studies

The mouse protocol (#2014-0078) was approved by the UTSW Animal IACUC (NIH OLAW Assurance Number A3472-01) and performed in accordance with UTSW IACUC guidelines. UTSW is fully accredited by the AAALAC. Animals are housed and maintained in accordance with the applicable portions of the Animal Welfare Act and the Guide for the Care and Use of Laboratory Animals. Veterinary care is under the direction of a full-time veterinarian boarded by the American College of Laboratory Animal Medicine. Mice were euthanized by Averin anesthesia followed by exsanguination.

### Kidney total RNA preparation for RNAseq

Kidneys were removed, and 20 mg of cortex was processed for RNA purification. All samples were stored with RNA later at -80 °C and thawed for RNA purification. Total RNA was purified using Direct-Zol^TM^ RNA Miniprep Plus (Zymo Research), and the concentration was measured with DS-11 FX+ (Denovix). Final RNA samples were submitted for sequencing to the Next Generation Sequencing Core of UTSW. Sequence analysis and results were performed by the University of Michigan Nephrotic Syndrome Study Network (NEPTUNE).

### Quantitative RT-PCR

Total RNAs were purified from each group of mouse cortices using Direct-Zol^TM^ RNA Miniprep Plus (Zymo research) and used for reverse transcription by SuperScript III First-Strand Synthesis System (Invitrogen). Quantitative PCR was carried out with specific primers using SYBR™ Green PCR Master Mix (Applied Biosystems) and StepOnePlus™ Real-Time PCR System (Applied Biosystems). For quantitative analysis, the samples were normalized to GAPDH (Glyceraldehyde 3-phosphate dehydrogenase) gene expression using the ΔΔCT value method. Primer sequences for specific genes are below. Primer sequences are given in **Table S19.**

### Measurement of glomerular stiffness

Glomerular stiffness (E, Young’s modulus) was measured using a microprobe indenter device as described previously using a microtensiometer (Kibron, Inc., Helsinki), a 3 -D micromanipulator with 160 nm step size (Eppendorf, Inc.) attached to a Leica inverted microscope(1, 2, 32). The absolute values for elastic modulus were measured as described(32). Indentations were <15 µm to avoid large strains that could damage the glomeruli.

### Imaging. Light microscopy of renal tissue

Formalin-fixed, paraffin-embedded kidney tissue was sectioned at 3 µm, and sections were stained with periodic acid-Schiff or picrosirius red stains using standard methods. Digital images were obtained with an Olympus BX-80 microscope equipped with a Q-Imaging Retiga 2000R imaging system. **Immunofluorescence staining.** Isolated glomeruli were processed and imaged as described(3). Glomeruli were fixed in 4% paraformaldehyde and attached to coverslips overnight, and permeabilized with 0.5% Triton X-100 in PBS. They were stained using standard procedures, and imaged using a Zeiss LSM 880 confocal microscope. **Confocal microscopy.** Confocal imaging was performed as previously described using a Zeiss LSM880 with Airyscan laser scanning microscope equipped with Plan-Apochromat 10x/0.3 NA, 20x/0.8 NA, 25x/0.8 NA and 63×/1.40 NA oil-immersion objective (ZEISS, Oberkochen, Germany). Fluorescence images were acquired using ZEN black 2.3 software with a 20x/0.8 NA or 63x/1.40 NA objective and Zeiss Immersion Oil 518F was used for the 63x/1.40 NA objective(3). Images or regions of interest (ROIs) were further processed with ZEN 2.6 (blue edition) software. **Electron microscopy.** Tissue samples were processed and imaged as described previously using standard methods(68). Images were acquired on a JEM-1400 Plus transmission electron microscope equipped with a LaB_6_ source operated at 120 kV using an AMT-BioSprint 16M CCD camera.

### Bulk RNA sequencing

The total input of 100ng RNA was used for the library preparation on each sample using Illumina TruSeq RNA Library Prep Kit v2 (RS-122-2001) according to the Manufacturer’s protocol. Illumina UMI barcodes were used to multiplex the libraries as per the Manufacturer’s instructions. The quality and quantity of each sequencing final library were assessed using the Agilent 2100 BioAnalyzer system (Agilent, CA, USA). A total of 10 nM picogreen measured library was pooled for 1.3pM loading on the sequencer. The pooled libraries were sequenced on the Illumina NexSeq platform in PE75 (75 bp paired end) run with the NextSeq reagent kit v2.5 for 75 cycles. Approximately 30-40 million sequencing reads were generated per sample for the transcriptome analysis.

Paired-end demultiplexed fastq files were generated using bcl2fastq2 (Illumina, v2.17), from NextSeq550 v2.5 reagents’ bcl files. Initial quality control was performed using FastQC v0.11.8 and multiqc v1.7. Fastq files were imported batch-wise, trimmed for adapter sequences, followed by quality trimming using CLC Genomics Workbench (CLC Bio, v23.0.3). The imported high-quality reads were mapped against gene regions and transcripts annotated by ENSEMBL v99 hg38 using the RNA-seq Analysis tool v2.7 (CLC Genomics Workbench); only matches to the reverse strand of the genes were accepted (using the strand-specific reverse option). Differential gene expression analysis between the sample groups was done using the Differential Expression for RNA-seq tool v2.8, where the normalization method was set as TMM. Differential expression between the groups was tested using a control group. Outliers were downweighed, and a filter on average expression for FDR correction was enabled.

### Gene expression QC, quantification, and DEG analysis

Fastq reads were aligned to the mouse genome (GENCODE Version M27), followed by a multi-step quality control process, which included assessments at the read, mapping, and dataset levels according to RNA sequencing quality control methods previously outlined(69). Gene-level expression quantification utilized HTSeq, and the resulting count data were normalized using Voom. Differential gene expression analysis was performed in R using the Limma package(70). The Benjamini & Hochberg method for controlling the false discovery rate was applied as a correction factor for the adjusted p-value. Statistically significant differentially expressed genes (DEGs) were determined based on an adjusted p-value threshold of less than 0.05. All heatmaps were created using the ComplexHeatmap R package (62). Upregulated and downregulated biological and cellular processes as well molecular function were performed using the Gene Ontology (GO) tool(36).

The Podocin: Nephrin ratio (PNR), a term introduced by Roger C. Wiggins, serves as a crucial marker for assessing podocyte hypertrophic stress by examining the differential mRNA expression of Podocin (NPHS2) and Nephrin (NPHS1). Under hypertrophic stress, podocytes exhibit an increase in NPHS2 and NPHS1 mRNA, with a disproportionately higher rise in NPHS2. This ratio has been shown to correlate with podocyte density, podocyte detachment, proteinuria, progressive glomerulosclerosis, and end-stage kidney disease (ESKD)(39). Elevated PNR indicates reduced podocyte density and increased detachment, both of which contribute to proteinuria and glomerulosclerosis, ultimately progressing to ESKD. Nishizono et al. provided robust evidence supporting the association between PNR and these pathological conditions and treatment effects, underscoring its potential as a diagnostic and prognostic tool in kidney disease(71).

### Identification of a TNF signature in Mouse RNAseq data

The TNF network reported in Mariani et al., 2023, which was generated from expert-curated interactions from NETPro annotations in the Genomatix Genome Analyzer database (Precigen Bioinformatics, Germany), was used to create the TNF activation score of 7-month-old mice(40). First, the mouse orthologs of the 272 genes from that paper were obtained from the Ensembl Biomart, yielding a TNF network of 205 mouse genes. Individual gene expression values were Z-transformed, and group mean gene expression was calculated. The TNF activation score for each group was the average Z-score of the 205 genes.

### Using human FSGS expression data at a single nuclear resolution to deconvolute 4-month-old mouse RNA-seq data

We identified 77 patients from the NEPTUNE cohort with a diagnosis of FSGS. Molecular data in the form of snRNA-seq were available for analysis and used in this study. Using expression data from 4-month-old mice, an Independent Component Analysis was performed to identify clusters of co-expressed genes in an unsupervised way. ICA is a matrix decomposition method that separates multifactorial signals into additive subcomponents. Each component could be driven by latent factors such as disease, cell type, and even noise. Twenty-five optimal modules were extracted using Python package stabilized-ica-2.0. The optimal module number was determined as described(72). We identified one module enriched for podocyte-specific genes. Because Alport nephropathy, like FSGS, is a primary podocytopathy, we used this module as a marker of podocyte injury (**Supplemental Figure S8**). Furthermore, we reasoned that because injury patterns are likely to be conserved across disease types and species, identifying the presence of these genes and their differential expression across all the time points (2, 4, and 7 months) will offer a temporal perspective on how these human data-enriched podocyte genes change over time. Their alteration with TUDCA therapy will provide clues to temporal changes in these genes and to the therapeutic potential of TUDCA or signals downstream of TUDCA.

## Supporting information

Yoon et al Supp Data

Yoon et al Supp Tables

## Author contributions

JY and ZL conducted experiments, JY, ZL, OA, and IM acquired data, JY, ZL, MA, CAA, OA, IM, SE, MK, JMH, VN, ASN, ANC, and RTM analyzed data, LAB and PAJ provided critical methods and reagents and advice, ASN, ANC and RTM wrote and edited the manuscript, all authors contributed to editing the manuscript.

## Acknowledgments

This work was supported by the National Institutes of Health grant numbers DK083592 (to LAB PAJ and RTM), DK139111, and R56DK139111 (RTM), the Charles and Jane Pak Center for Mineral Metabolism and Clinical Research (RTM, ANC), and the NEPTUNE consortium (ASN, MK, VN, SE, MA). ASN acknowledges support of Intramural grants from the University of Michigan O’Brien Kidney Translational Core Center P30 DK081943, Taubman Institute and an extramural grant from the National Institute of Health K23 DK 125529. RTM appreciates support from the VANTHCS Research Service and COS, Jeff Hastings.

The <u>Nephrotic Syndrome Study Network</u> (NEPTUNE) is part of the <u>Rare Diseases Clinical Research Network</u> (RDCRN), which is funded by the National Institutes of Health (NIH) and led by the <u>National Center for Advancing</u> <u>Translational Sciences</u> (NCATS) through its <u>Division of Rare Diseases Research Innovation</u> (DRDRI). NEPTUNE is funded under grant number U54DK083912 as a collaboration between NCATS and the <u>National</u> <u>Institute of Diabetes and Digestive and Kidney Diseases</u> (NIDDK). Additional funding and/or programmatic support is provided by the University of Michigan, NephCure Kidney International, Alport Syndrome Foundation, and the Halpin Foundation. RDCRN consortia are supported by the RDCRN Data Management and Coordinating Center (DMCC), funded by NCATS and the <u>National Institute of Neurological Disorders and Stroke</u> (NINDS) under U2CTR002818.

