## Supplementary material for "Glomerular Elastic Modulus and Gene Expression Patterns Define Two Phases of Alport Nephropathy": Yoon et al Supp Data

**Suppemental Figure S1**

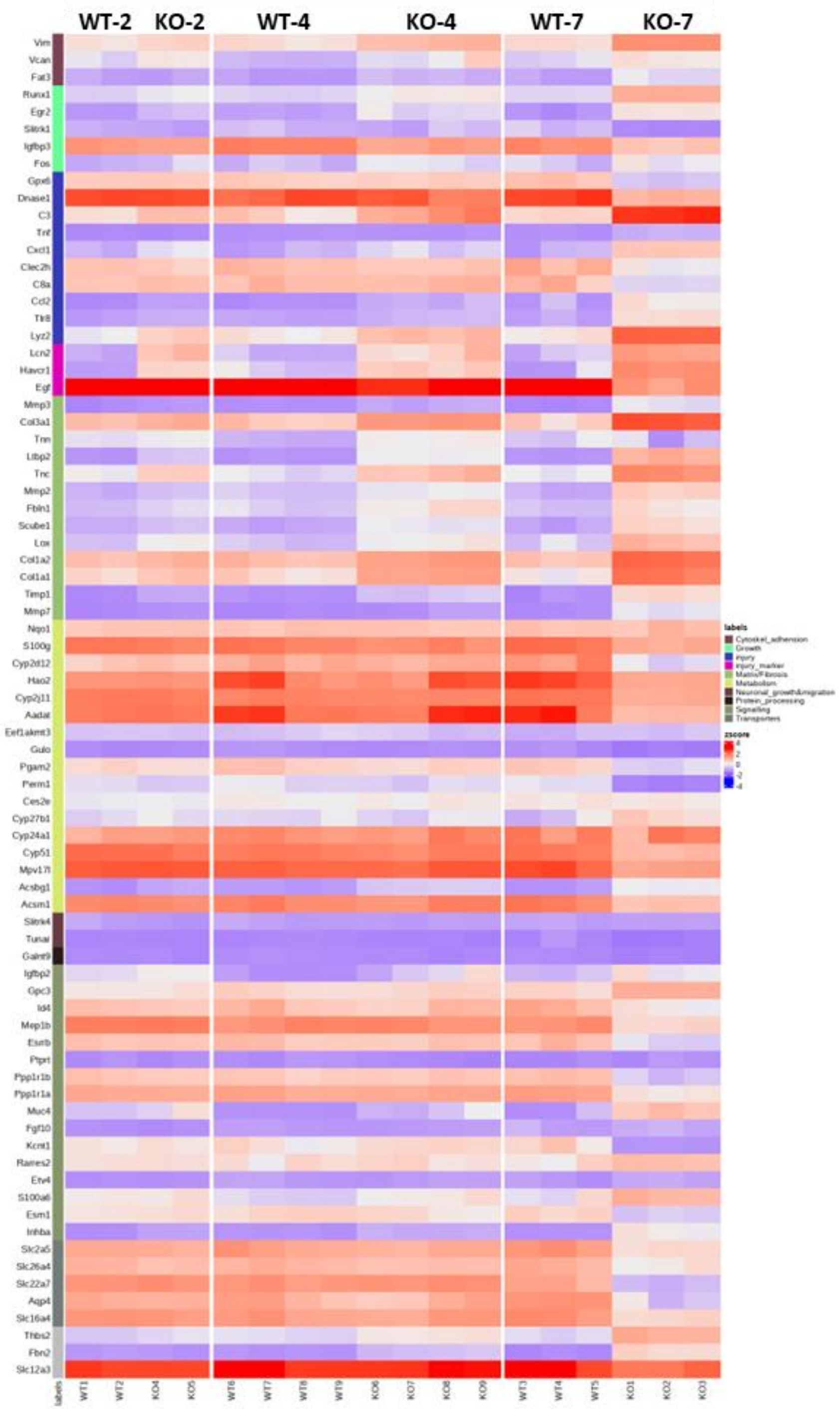


**Supplemental Figure S1**. **Heatmap of genes in the lists of DEGs from WT and Col4α3$^{-/-}$ (KO) mice at 2, 4, and 7 months.** The heatmap shows the samples across the top, WT and Col4α3$^{-/-}$ (KO) at 2 months (2 samples each), WT and KO 4 months (four samples each), and WT and KO at 7 months each with three samples each. Comparisons are to age-matched WT controls. Gene names are grouped on the left side by function. Red represents increased and blue represents decreased relative expression (Complex Heatmap R Package).

### Supplemental Figure S2

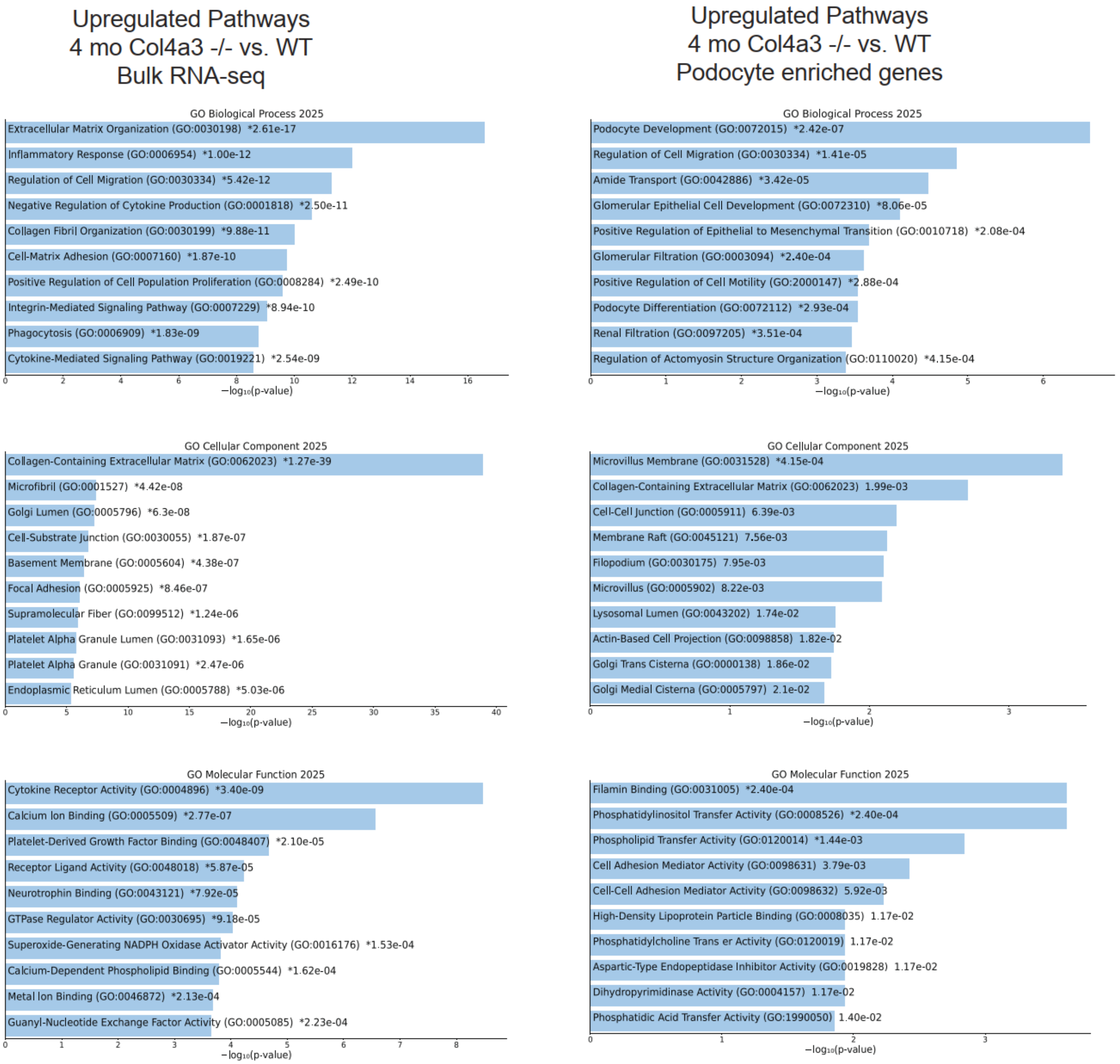


**Figure S2:** Bar chart of top upregulated terms from 4-month-old Col4α3-/- mice vs. 4-month old WT mice. Terms are categorized based on their Biological processes, Cellular components and Molecular function using the GO 2025 gene library. The top 10 enriched terms for the input gene set are displayed based on the -log10(p-value) with actual p-value adjacent to it. * Denotes that enrichment term also reached FDR adjusted p-value. Note: A negative regulation of any process in a query of upregulated pathways or vice versa denotes that the input genes set are over represented in a process that bears the opposite direction to that of the query. The left panel uses the entire bulk RNA-seq data, and the right panel shows enrichment from the podocyte-enriched gene set.

**Supplemental Figure S3A**

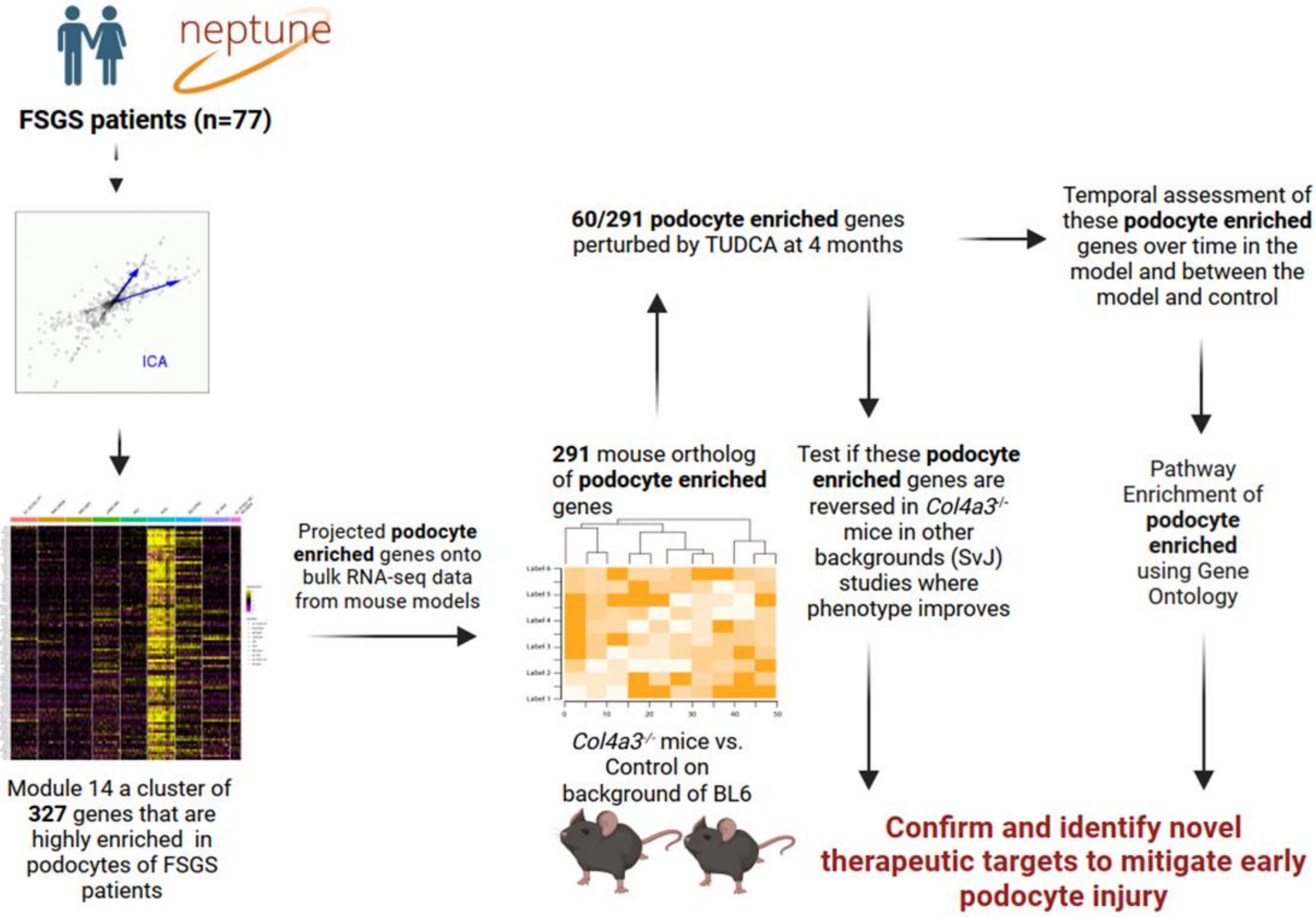


**Figure S3A:** Analytic framework to obtain a podocyte-enriched gene set from humans that was used to deconvolute mice bulk-RNA seq data.

**Supplemental Figure S3B**

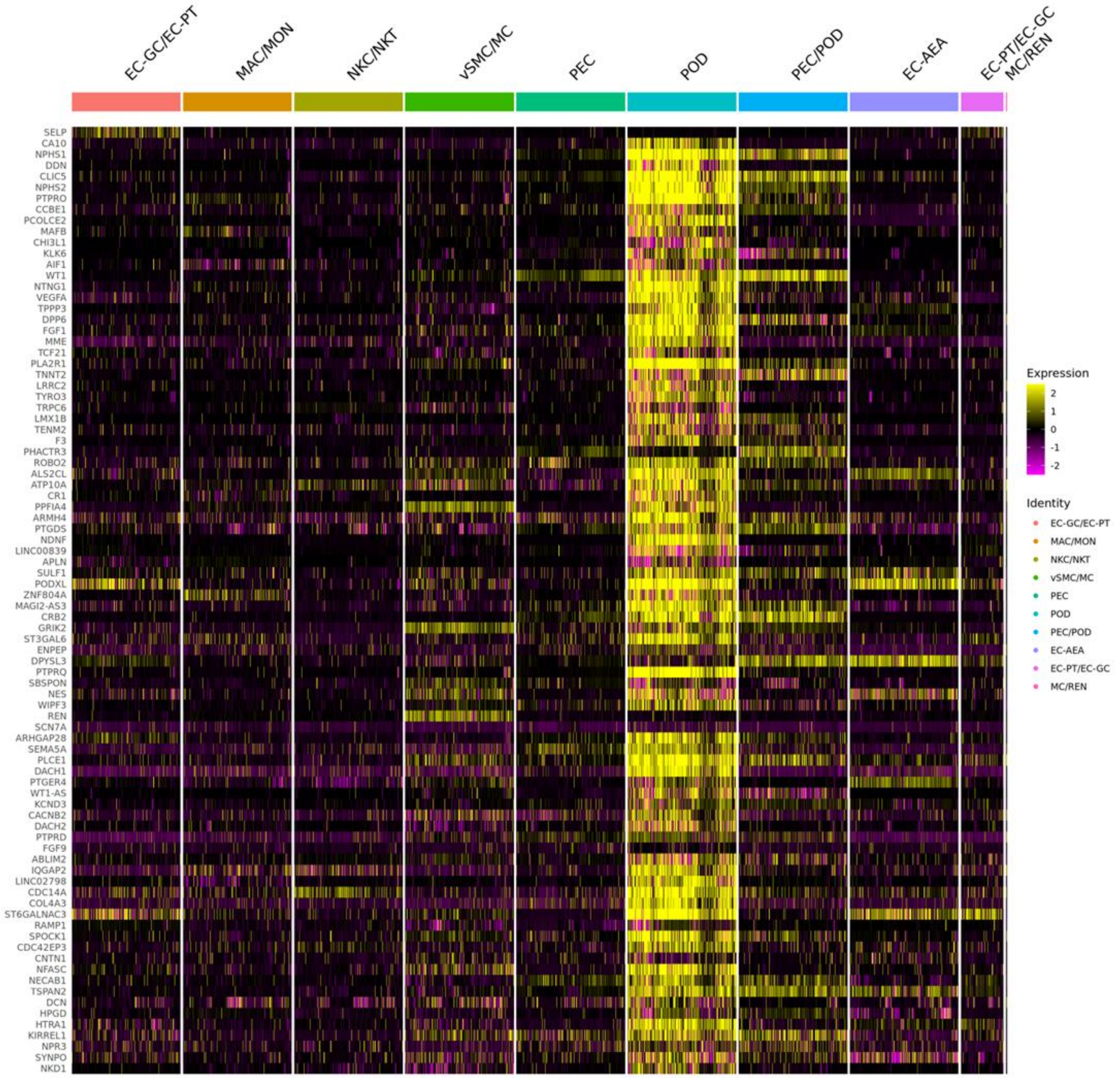


**Figure S3B:** The heatmap reflects the gene module obtained from bulk-RNA seq data of micro-dissected glomeruli from proteinuric patients in the NEPTUNE consortium. The clustering was performed using Independent Component Analysis.

#### Supplemental Figure S4

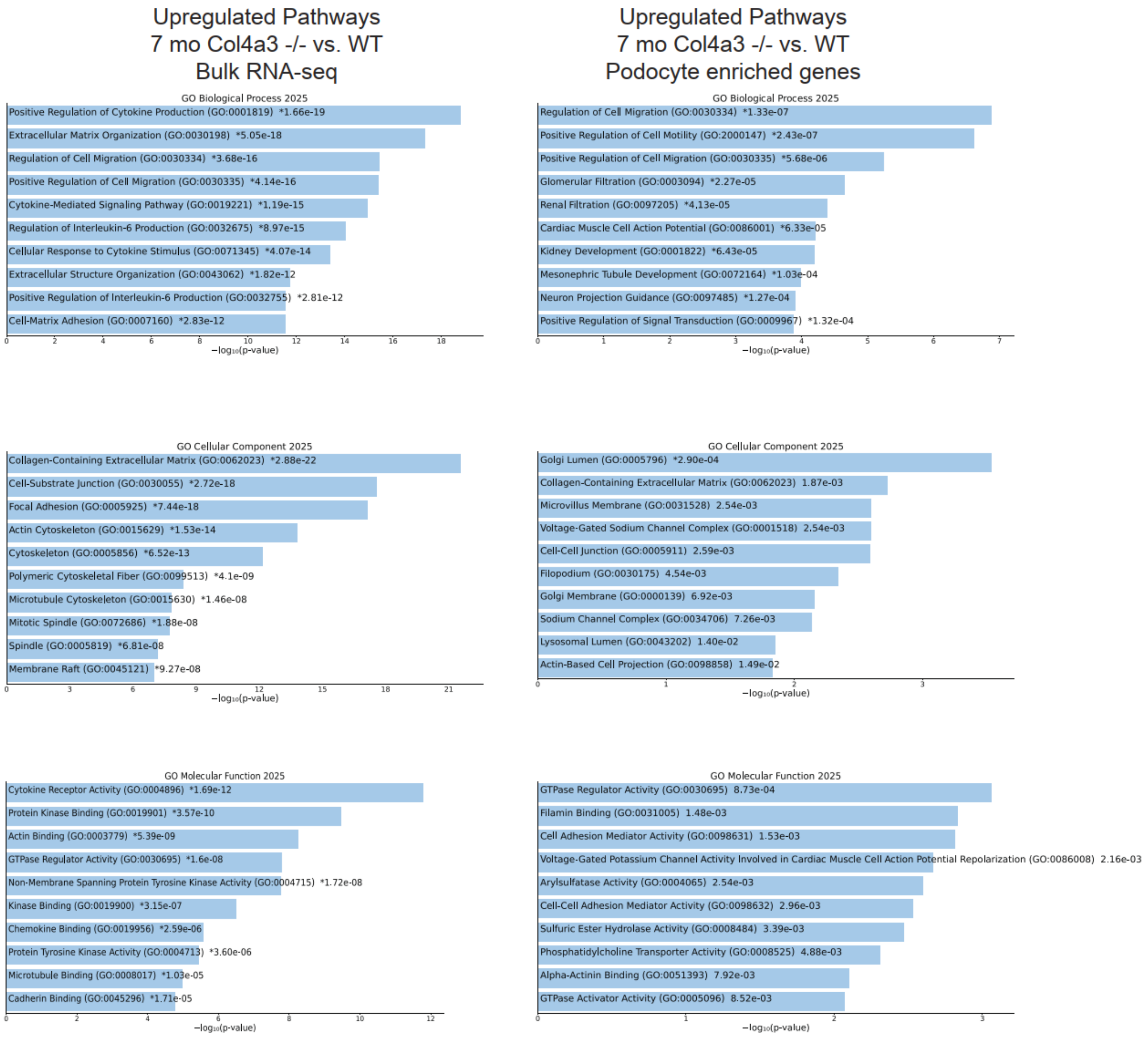


**Figure S4:** Bar chart of top upregulated terms from 7-month-old Col4α3-/- mice vs. 7-month old WT mice. Terms are categorized based on their Biological processes, Cellular components and Molecular function using the GO 2025 gene library. The top 10 enriched terms for the input gene set are displayed based on the -log10(p-value) with actual p-value adjacent to it. * Denotes that enrichment term also reached FDR adjusted p-value. Note: A negative regulation of any process in a query of upregulated pathways or vice versa denotes that the input genes set are over represented in a process that bears the opposite direction to that of the query. The left panel uses the entire bulk RNA-seq data, and the right panel shows enrichment from the podocyte-enriched gene set.

#### Supplemental Figure S5

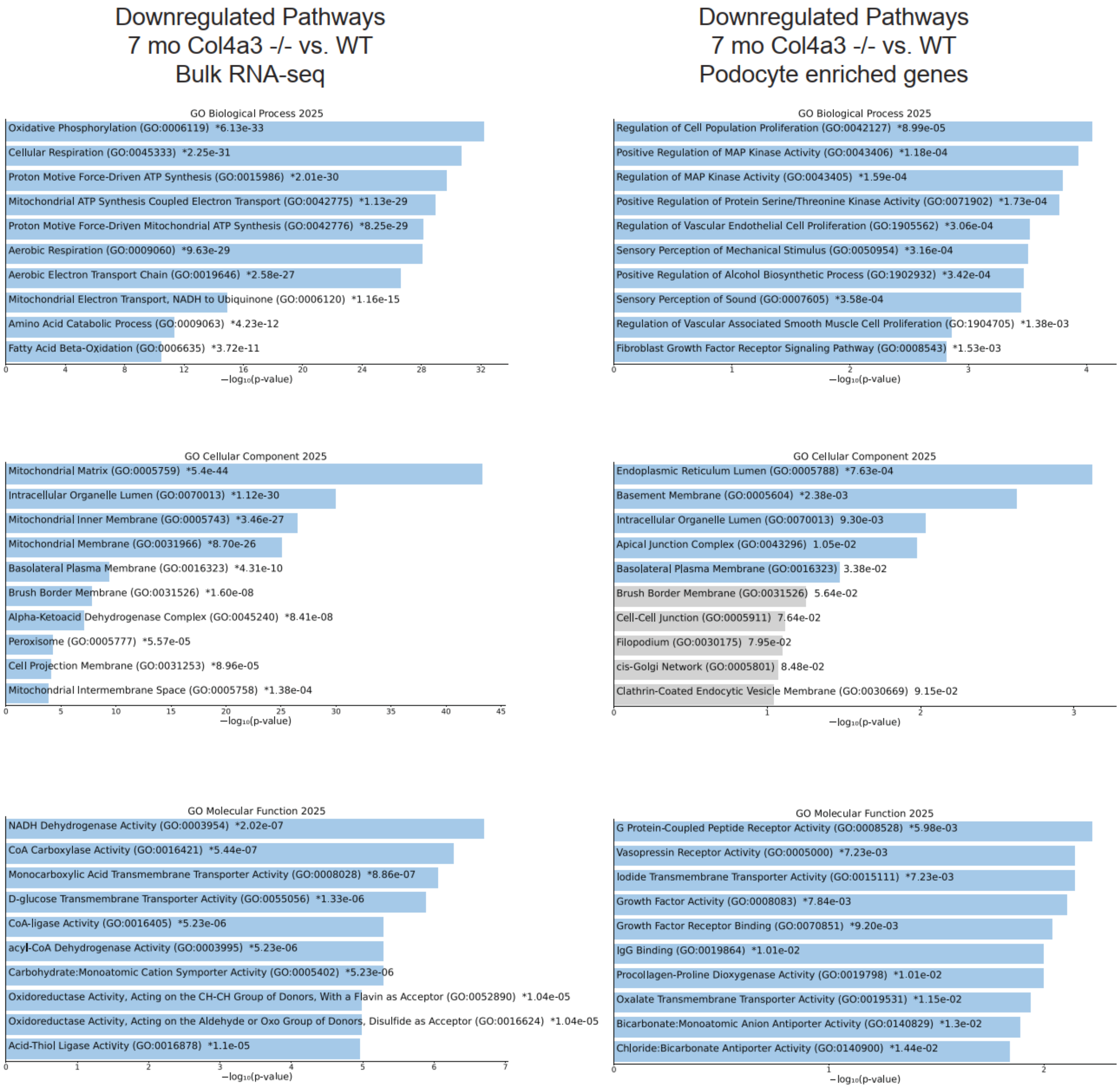


**Figure S5B:** Bar chart of top downregulated terms from 7-month-old Col4α3-/- mice vs. 7-month old WT mice. Terms are categorized based on their Biological processes, Cellular components and Molecular function using the GO 2025 gene library. The top 10 enriched terms for the input gene set are displayed based on the -log10(p-value) with actual p-value adjacent to it. * Denotes that enrichment term also reached FDR adjusted p-value. Note: A positive regulation of any process in a query of downregulated pathways or vice versa denotes that the input genes set are over represented in a process that bears the opposite direction to that of the query. The left panel uses the entire bulk RNA-seq data, and the right panel shows enrichment from the podocyte-enriched gene set.

**Supplemental Figure S6**

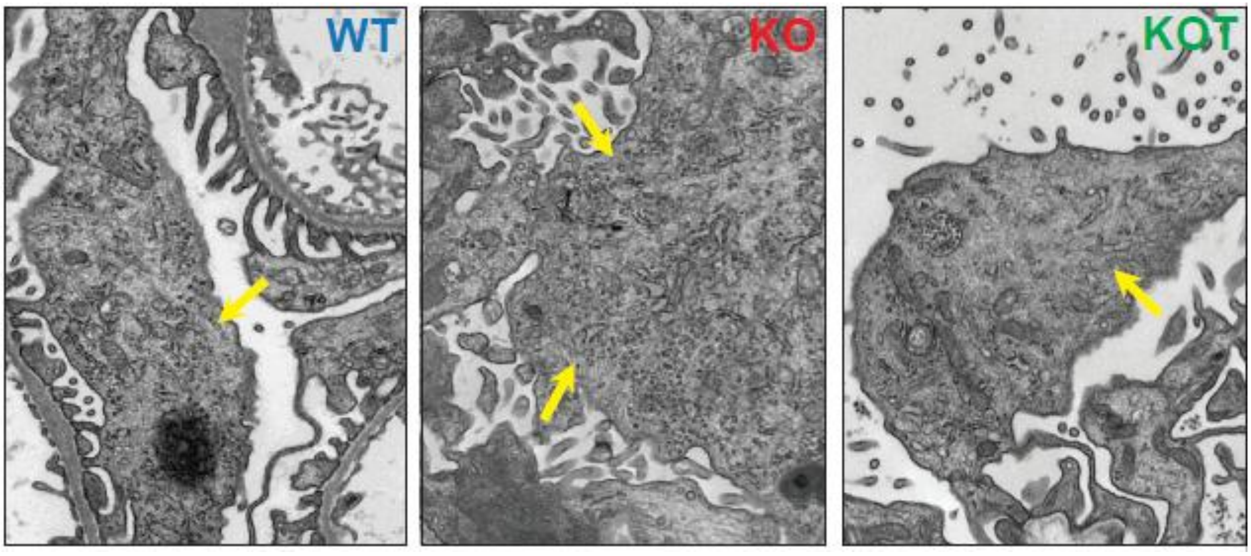


**Figure S6:** Representative TEM images of podocyte cell bodies at 4 Mo from WT, *Col4α3*$^{-/-}$, TUDCA-treated *Col4α3*$^{-/-}$ mice. The podocyte cell bodies of the *Col4α3*$^{-/-}$ mice are hypertrophic and the ER is prominent with increased density, increased cross-sectional size and complexity of shape, and more obvious ribosomes compared to the WT cell. The state of the TUDCA-treated *Col4α3*$^{-/-}$ podocyte (KOT) appears intermediate between WT and *Col4α3*$^{-/-}$ phenotypes. Yellow arrows mark ER.

**Supplemental Figure S7**

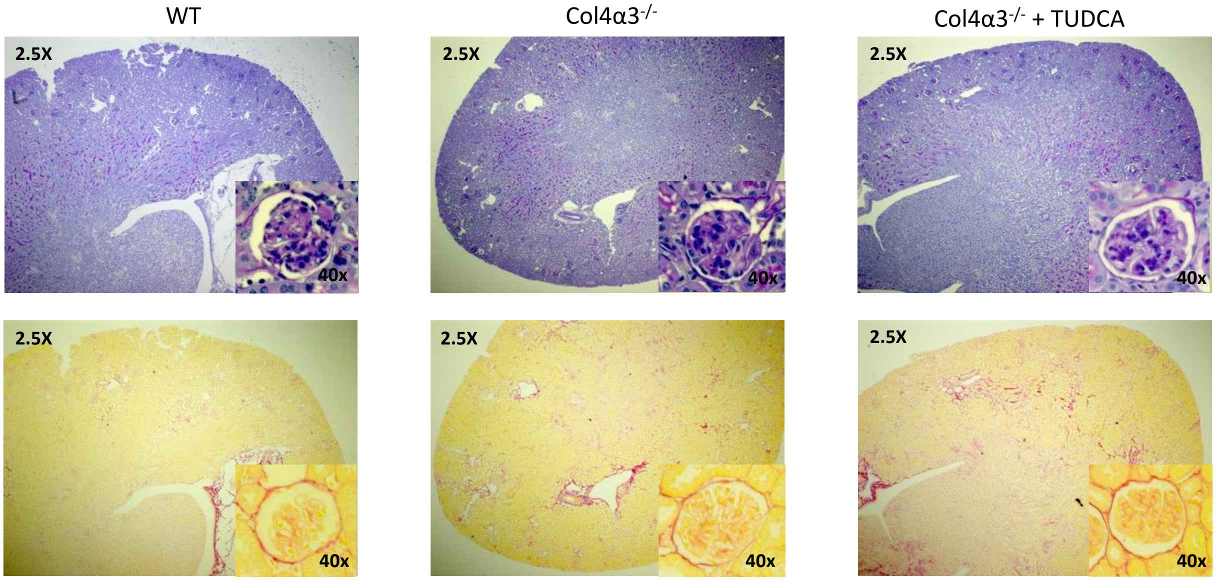


**Figure S7. Minimal renal pathology at low magnification in 4 month WT, *Col4α3*$^{-/-}$, and *Col4α3*$^{-/-}$ mice treated with TUDCA.** Mice were treated with TUDCA from weaning (approximately 4 weeks, with 0.8% TUDCA mouse chow, 3 months of TUDCA exposure). Sections were stained with PAS and picrosirius red as described for Figure 3. Insets show representative glomeruli from each group. The differences among the groups are minimal at this magnification.

**Supplemental Figure S8**

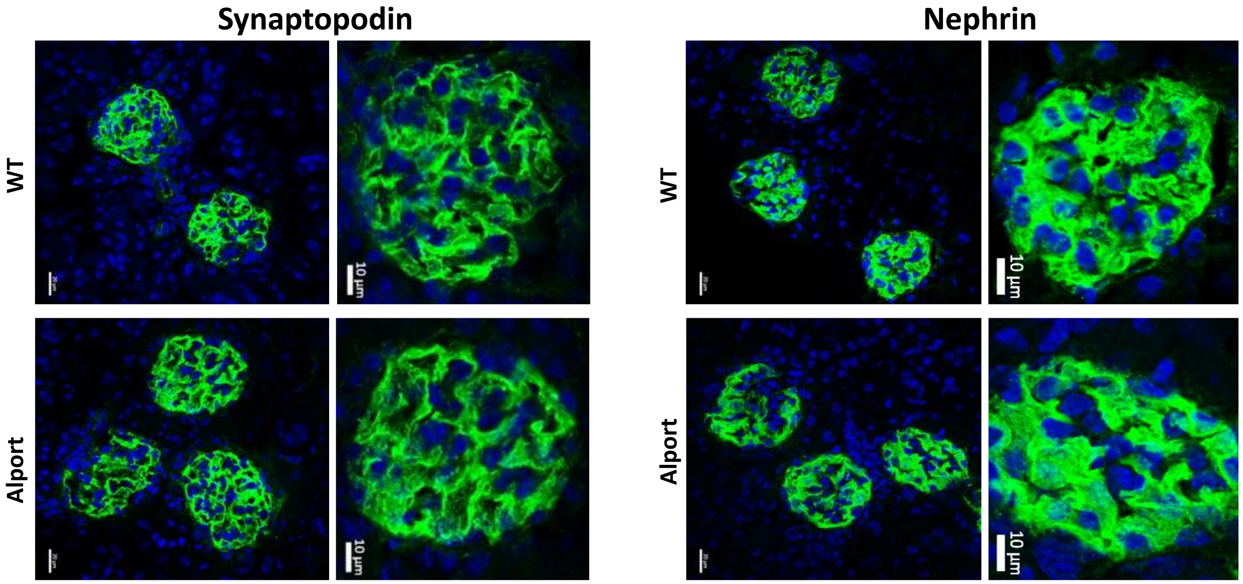


**Figure S8**: Confocal (Z-stack) images of isolated 3 Month WT or *Col4a3*$^{-/-}$ Glomeruli Stained for Synaptopodin or Nephrin. At 3 months, the structure of the *Col4α3*$^{-/-}$ and WT glomeruli appears similar indicating minimal if any significant structural damage at this magnification in the *Col4α*$^{-/-}$ glomeruli.

#### Supplemental Figure 9

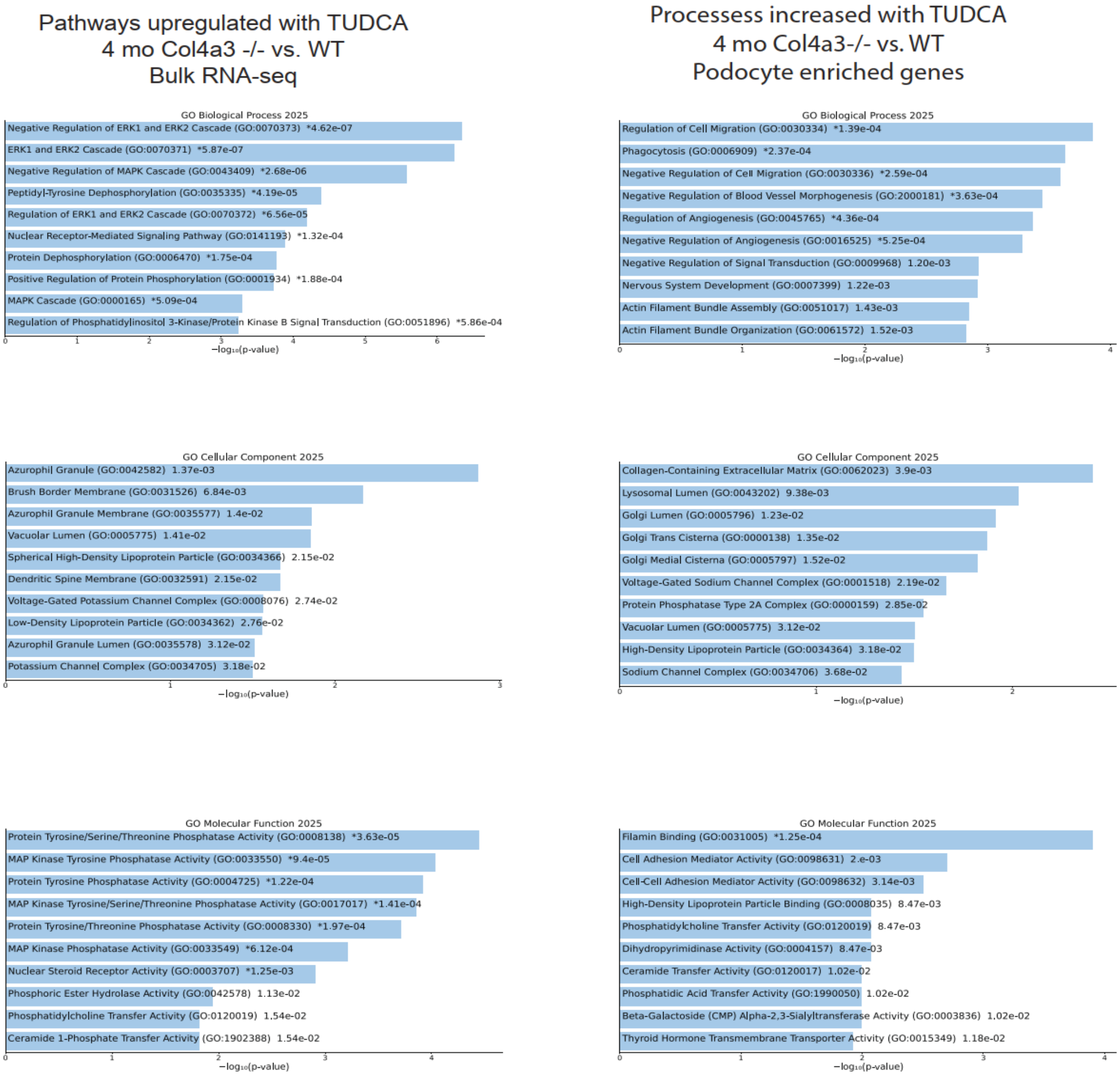


**Figure S9:** Bar chart of top enriched terms from GO, Biological processes, Cellular component and Molecular function 2025 gene library. The top 10 enriched genes for the input gene set are displayed based on -log10(p-value) with actual p-value next term. The term at the top has the the most significant overlap with the input gene set. * Denotes that enrichment term also reached FDR adjusted p-value. Note: A negative regulation of any processess within an upregulated pathways or vice versa denotes that the input gene sets are over-representated in a processes that bears the opposite direction to that of the pathway.

#### Supplemental Figure 10

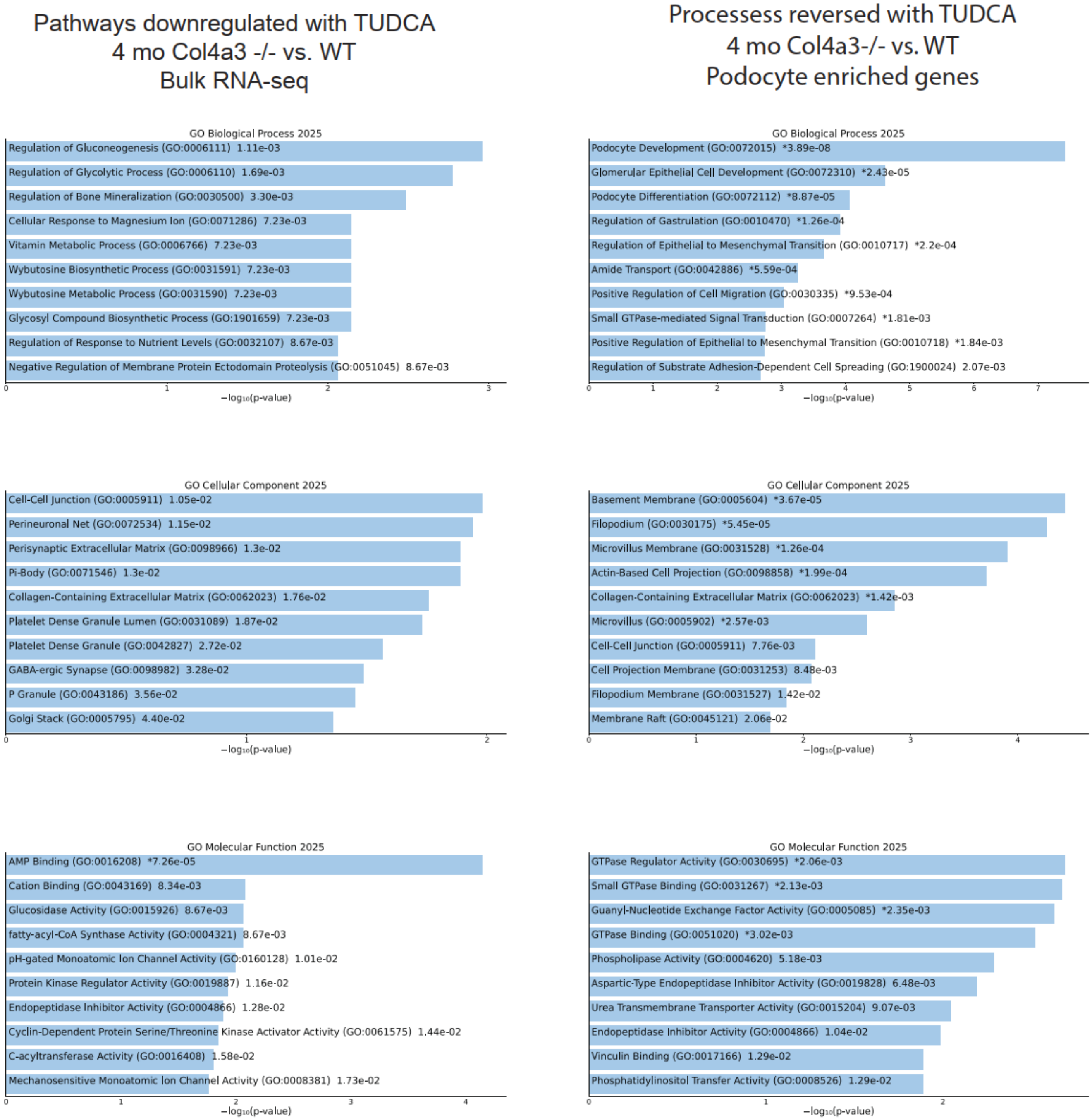


**Figure S10:** Bar chart of top downregulated terms from 4-month-old Col4α3-/- mice treated with TUDCA vs. no TUDCA. Terms are categorized based on their Biological processes, Cellular components and Molecular function using the GO 2025 gene library. The top 10 enriched terms for the input gene set are displayed based on the -log10(p-value) with actual p-value adjacent to it. * Denotes that enrichment term also reached FDR adjusted p-value. Note: A negative regulation of any process in a query of upregulated pathways or vice versa denotes that the input genes set are over represented in a process that bears the opposite direction to that of the query. The left panel uses the entire bulk RNA-seq data, and the right panel shows enrichment from the podocyte-enriched gene set.

**Supplemental Figure S11**

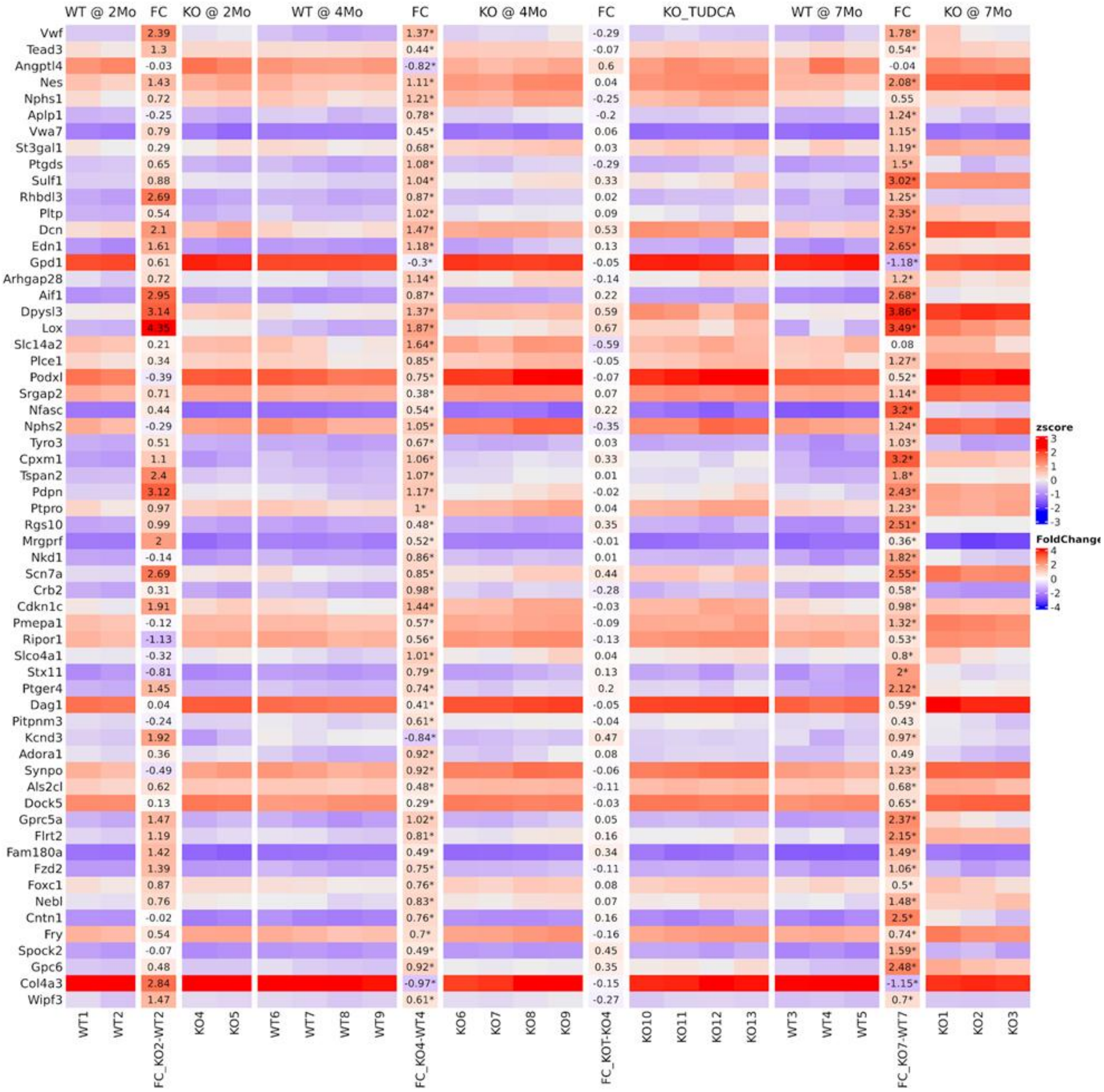


**Figure S11**: Heatmap of the expression of 60 podocyte-enriched TUDCA responsive genes in WT 2 month, *Col4α3*$^{-/-}$ 2-month, WT 4 month, *Col4α3*$^{-/-}$ 4 month, *Col4α3*$^{-/-}$ +TUDCA 4 months, WT 7 months, and *Col4α3*$^{-/-}$ 7 month mice. Mouse genotypes are shown along the top, with each column representing an individual mouse. Genes are shown on the left. The vertical bars separating heatmaps show the log2 fold change between groups to the left and right of each bar and colors represent increased (Red) and decreased (Blue) relative expression compared to age-matched WT controls. The third column shows the comparison between 4 Mo *Col4α3*$^{-/-}$ mice and 4 Mo *Col4α3*$^{-/-}$ mice treated with TUDCA. In this comparison, TUDCA caused sign reversal in 26 genes and reduced the magnitude of change in re remainder compared to *Col4α3*$^{-/-}$ mice. These values correspond to those shown in Table S6. Genes enriched in podocytes were obtained from 77 human FSGS samples from the NEPTUNE consortium.

**Supplemental Figure S12**

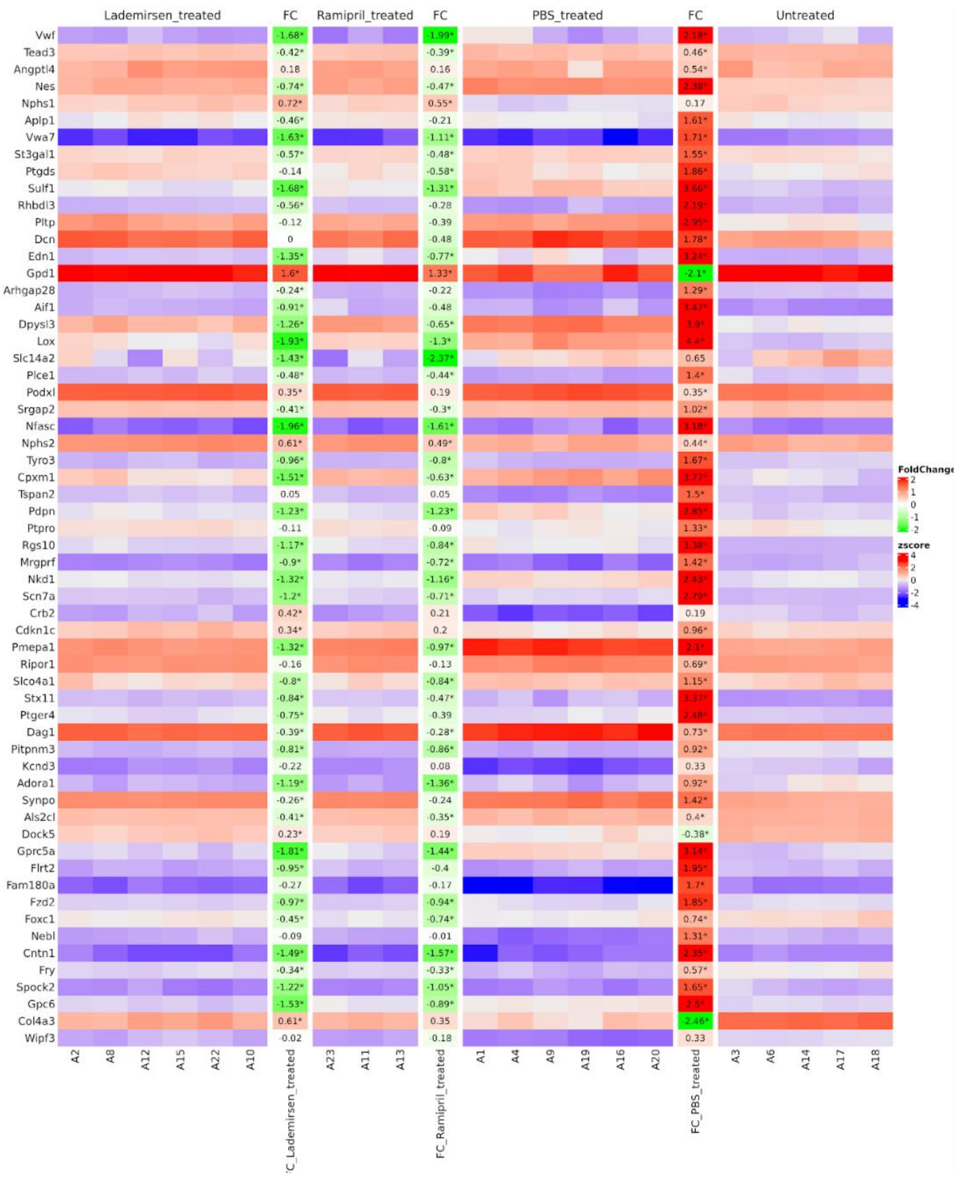


**Figure S12:** Behavior of the 60 TUDCA-responsive podocyte-enriched genes in *Col4α3*$^{-/-}$ mice applied to a database from F1 hybrid Svj126 X Bl6 *Col4α3*$^{-/-}$ mice treated with Lademirsen (anti miR21), Ramipril, PBS, or WT (Ref 485, Rubel et al. Cells. 2022; 11(4) (Reference 45). The treatments are shown across the top. Each column represents a single mouse. Untreated = WT, Lademirsen - treated = Col4a3$^{-/-}$ + anti-miR-21, Ramipril-treated = Col4a3$^{-/-}$ + Ramipril, and PBS-treated = Col4a3$^{-/-}$ + PBS. The vertical bars separating heatmaps show the log2 fold change between groups and colors represent increased (Red) and decreased (Green) relative to untreated WT with notations for the comparisons. The left margin identifies the genes which and they are the same as those shown in Figure S11 and Table S6. The samples from Rubel et al were obtained at 15 Weeks, probably a later stage of disease than our 4 Month samples. * denotes significance <0.05 (Complex Heatmap R Package).

#### Supplemental Figure 13

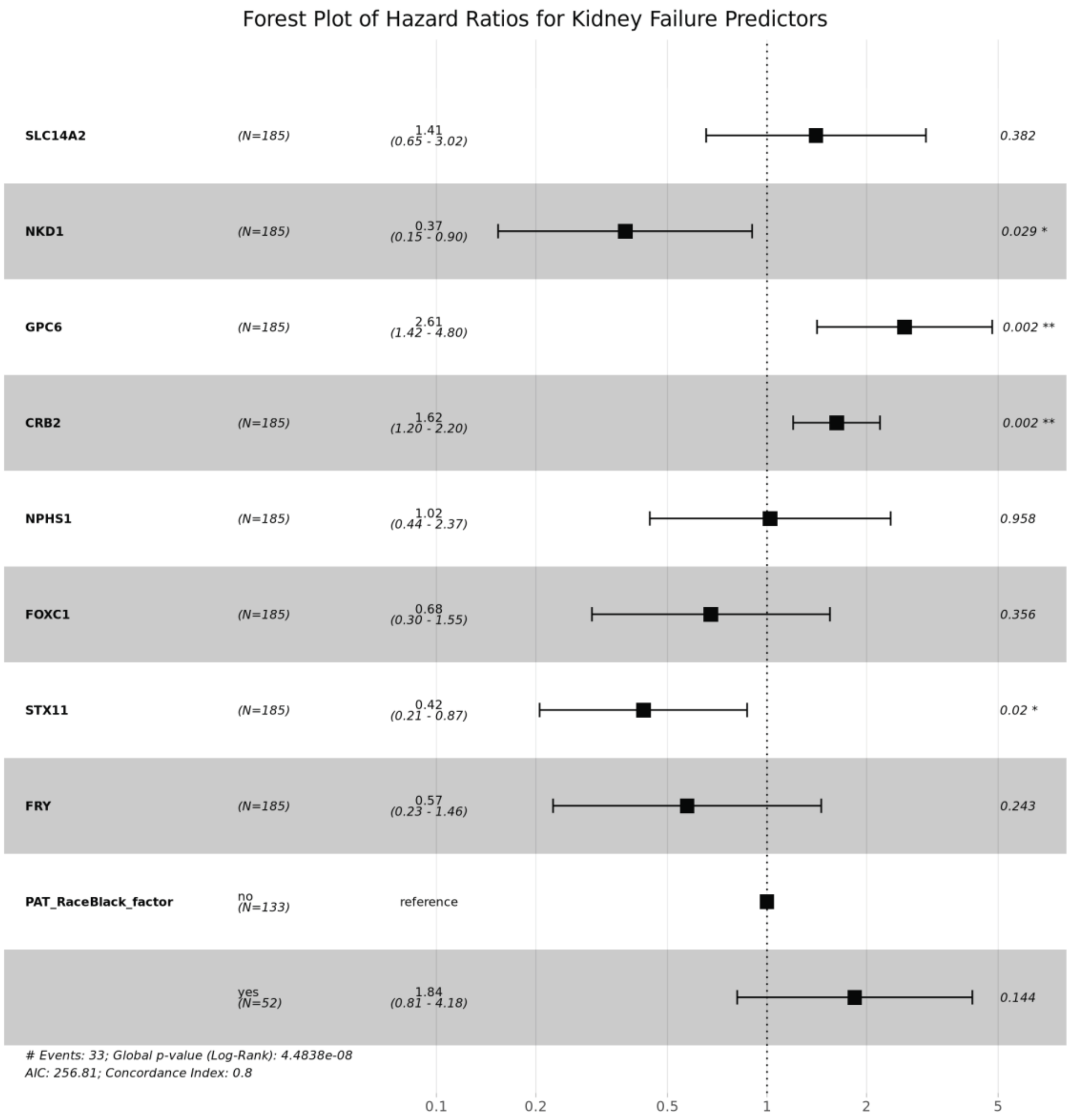


**Figure S13**: Forest plot of genes associated with the composite of ESRD or a 40% decline in eGFR from baseline (ESRD40). The model was adjusted for age, gender and african american race.

**Supplemental Figure S14**

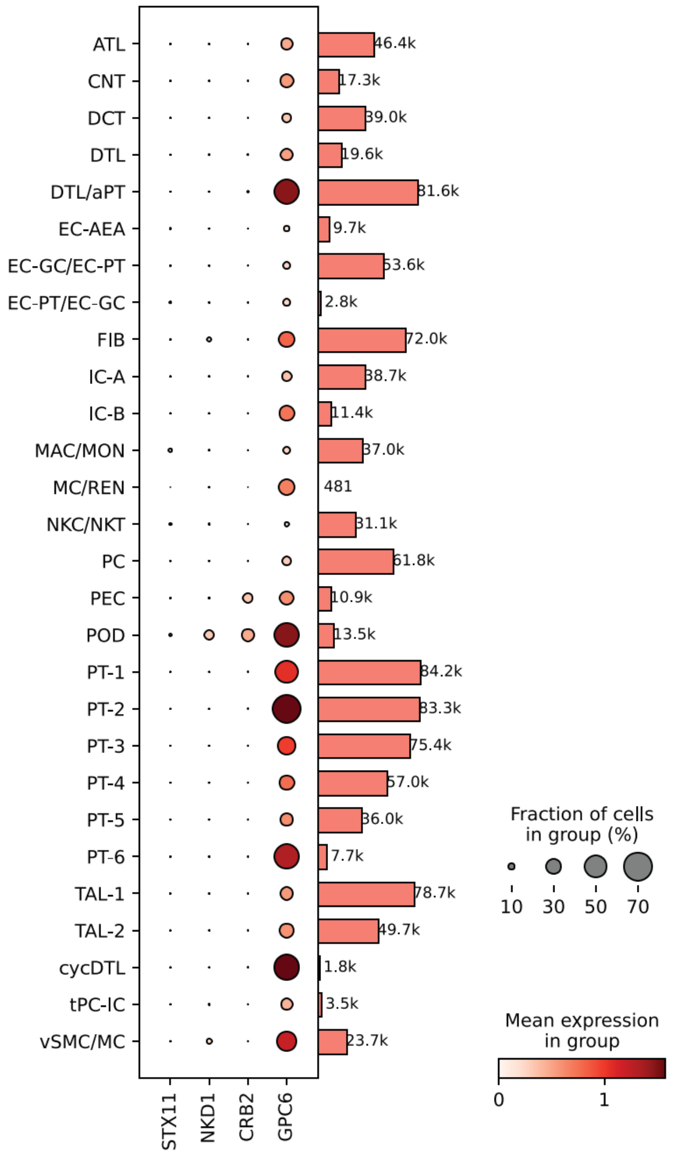


**Supplemental Figure S14. Dot plot showing expression of *STX11* (Syntaxin-11), *NKD1* (NKD Inhibitor of Wnt Signaling Pathway 1), *CRB2* (Crumbs Cell Polarity Complex Component 2), and *GPC6* (Glypican 6) in cells of kidney tissue obtained from patients in the NEPTUNE study who underwent single nuclear RNA sequencing.** Three of the 4 genes (*NKD1, CRB2, GPC6*) associated with ESRD40 are expressed higher in podocytes than other renal cell types undergoing single nuclear RNA- sequencing. This data provides parallel validation that the deconvolution method to obtain podocyte enriched genes was an effective approach. *STX11* gene expression was minimal in podocytes and all other cell types. Cell designations are as follows: ATL (Ascending Thin Limb), CNT (Connecting Tubule), DCT (Distal Convoluted Tubule), DTL (Descending Thin Limb), C-DTL- cycling DTL cells, DTL_aPT (Descending thin limb_adaptive proximal tubule), EC1 (Arteriolar Endothelial Cells), EC2 (Glomerular Endothelial Cells), EC3 (Peritubular Capillaries), FIB (Fibroblasts), IC A/B (Intercalated Cells A and B, MAC (Macrophages), MC (Mesangial Cells), MON (Monocytes), NKC (Natural Killer Cells), NKT (Natural Killer T Cells), PC (Principal Cells), PEC (Parietal epithelial cells), POD (Podocytes), PT/DTL (Proximal Tubule/Descending Thin Limb), PT-1/2/3/4/5/6 (Proximal Tubule Sub-clusters 1, 2, 3, 4,5,6), REN (Renin secreting cells) TAL1/2 (Thick et al. 1 or 2 Sub-cluster), tDCT_CNT (Transitional cells of DCT with CNT), tPC_IC (Transitional Principal and Intercalated Cells), tDAL_DCT (Transitional TAL and DCT cells), vSMC (Vascular Smooth Muscle Cells).

| | Treatment | Global Sclerosis (%) | Segmental Sclerosis (%) | Hypo-perfusion (%) | Global Proliferation (%) | Segmental Proliferation (%) | Bowman's Space Distension (%) | Not Classifiable (%) |
|---|---|---|---|---|---|---|---|---|
| 4 months | Control | 0.0 | 0.0 | 0.0 | 2.6 | 2.6 | 0.0 | 0.0 |
| | KO | 0.0 | 0.4 | 0.8 | 7.5 | 5.7 | 0.0 | 1.1 |
| | KO TUDCA | 0.0 | 0.0 | 0.2 | 3.9 | 11.0 | 0.0 | 3.2 |
| 7 months | Control | 0.0 | 0.0 | 2.8 | 1.4 | 11.9 | 0.5 | 0.0 |
| | KO | 1.9 | 1.2 | 6.2 | 53.1 | 25.9 | 1.2 | 1.2 |
| | KO TUDCA | 0.0 | 0.0 | 6.8 | 14.1 | 21.1 | 0.9 | 3.3 |

**Supplemental Table S11. Morphometric analysis of glomerular lesions in light microscopic sections.** Results are presented as percentage of all glomeruli counted in the sample demonstrating the indicated lesion. Lesions classified include global and segmental sclerosis, glomerular hypoperfusion, global and segmental proliferation (both endo- and extra-capillary proliferation and including both epithelial proliferation and inflammatory cell infiltration), distension of Bowman's space with no changes of the glomerular tuft. Overall, glomerular lesions were focal in KO and KO-TUDCA mice, and rare in Control mice at 4 months, and typically consisted of global or segmental proliferation. At 7 months, KO mice showed widespread global proliferative lesions, involving greater than half of all glomeruli observed in the sample. In contrast, KO-TUDCA mice showed predominantly segmental proliferation and over half of the glomeruli exhibiting no lesion.

| | Treatment | Foot Process Effacement | Foot Process Irregularity | Podocyte ER Prominence | GBM “Knobs” (redundancy) | Capillary Dilation |
|---|---|---|---|---|---|---|
| 4 months | WT | 0 | 0 | 1 | 0 | 0 |
| | KO | 1 | 2 | 2 | 3 | 1 |
| | KO TUDCA | 1 | 2 | 2 | 2 | 1 |
| 7 months | WT | 0 | 0 | 0 | 0 | 1 |
| | KO | 3 | 3 | 2 | 3 | 2 |
| | KO TUDCA | 1 | 1.5 | 1 | 1.5 | 1 |

**Supplemental Table S12. Semi-quantitative morphometric analysis of key features of glomerular injury from transmission electron microscopy images.** Each feature was scored on the scale: 0 (no injury or less than 10% involvement), 1 (mild injury or 10 to 25% involvement), 2 (moderate injury or 25 to 50% involvement), and 3 (severe injury or greater than 50% involvement). Each of the features is present at some level in KO mice, but is less severe in TUDCA-treated mice at 7 months. For WT and KO mice, n=1, and for KO-TUDCA mice, n=2 (result presented is average of both specimens). “ER” = endoplasmic reticulum; “GBM” = glomerular basement membrane.
